# scDecorr - Feature decorrelation representation learning with domain adaptation enables self-supervised alignment of multiple single-cell experiments

**DOI:** 10.1101/2024.05.17.594763

**Authors:** Ritabrata Sanyal, Yang Xu, Hyojin Kim, Rafael Kramann, Sikander Hayat

## Abstract

Single-cell RNA sequencing (scRNA-seq) has revolutionized our understanding of cellular heterogeneity in complex biological systems. However, analyzing and integrating scRNA-seq data poses unique computational challenges due to sparsity, high variability, and technical batch effects. Here, we propose a novel framework called scDecorr for robust representation learning and data integration for scRNA-seq analysis. Our approach leverages the idea of feature decorrelation-based self-supervised learning (SSL) to obtain efficient low-dimensional representations of individual cells without relying on negative samples. By maximizing similarity among distorted embeddings while decorrelating their components, scDecorr captures the biological signature while eliminating technical noise. Furthermore, scDecorr incorporates unsupervised domain adaptation to bridge the gap between batches with different probability distributions, enabling effective integration of scRNA-seq data from diverse sources. Our framework achieves domain-invariant representations by learning cell embeddings independently across domains and employing domain-specific batch normalization. We evaluate scDecorr on a variety of single cell datasets and demonstrate its ability to integrate batches without losing the inherent biological variance, thereby facilitating optimal clustering. The representations generated by scDecorr also exhibit robustness in label transfer tasks, allowing for effective transfer of cell-type labels from reference to query datasets. Overall, scDecorr offers a powerful tool for efficient analysis and integration of large and complex scRNA-seq datasets, advancing our understanding of cellular processes and disease mechanisms. The code is available here https://github.com/hayatlab/scdecorr.

## 1 Introduction

Single-cell RNA sequencing (scRNA-seq) has emerged as a powerful tool to study cellular heterogeneity in complex biological systems[1, 2, 3] by enabling the measurement of gene expression at the individual cell level. However, scRNA-seq data presents unique computational challenges, including sparsity and high variability, which can adversely affect the accuracy of cell clustering and detection of rare cell populations. In this regard, obtaining a reliable low-dimensional representation for each cell that preserves the biological signature while eliminating technical noise is a crucial step in scRNA-seq data analysis[4, 5]. Furthermore, integrating scRNA-seq data from different sources, batches and platforms poses a significant challenge due to technical batch effects, which can confound the inherent biological variations[6, 7, 8]. These confounding factors can also influence downstream data analysis and interpretation. Various computational tools and pipelines, including Seurat[9], Harmony[10], MNN-Correct[11], scVI[12] have been developed to address these challenges. With the increasing availability of large and complex scRNA-seq datasets, there is a pressing need for accurate and efficient methods to analyze and integrate these data to increase our understanding of cellular processes and disease mechanisms.

The task of identifying an efficient low-dimensional, cell class discriminative, batch-invariant representations for single cells can be concisely defined as follows– The representation learning algorithm is required to map the gene expression profile onto a low-dimensional embedding, such that functionally similar cells are in close proximity while maintaining a distance between dissimilar cells. Additionally, the model must be resistant to batch effects, such that gene expression profiles belonging to the same cell type, regardless of their batch origin, are mapped to identical locations in the lower-dimensional space.

Computational tools and pipelines such as Seurat[9], Harmony[10], and scVI[12] have been developed to address this task. Recently, self-supervised learning (SSL) has gained a lot of traction for learning robust representations and solving various pretext tasks in multiple areas[13, 14, 15, 16, 17], including computer vision and natural language professing (NLP). In particular, contrastive learning (CL), a branch of SSL has emerged as a promising approach for finding robust low-dimensional single-cell representations[18, 19, 20, 21, 22]. CL enables representation learning by distinguishing positive pairs from negative pairs, where positive pairs consist of semantically similar transcriptional profiles and negative pairs consist of dissimilar ones. CL-based techniques construct positive cell pairs and negative cell pairs with the aim of concentrating positive pairs and separating negative pairs using a contrastive loss[23]. There are primarily two main strategies for constructing these pairs. The first strategy[21, 20, 24] involves computing inter-batch mutual nearest neighbors and intra-batch nearest neighbors for constructing positive pairs. The negative pairs are usually constructed by random sampling. This strategy introduces an overhead of computing the nearest neighbors in every iteration of the training run. The second strategy generates positive pairs of an anchor cell gene profile via random augmentations and constructs negative pairs by random sampling from a memory bank [25, 26] or by using the current mini-batch[22, 13]. These methods typically require a large number of negative samples to generate high-quality representations and thus have a high memory footprint. Moreover, these techniques usually also suffer from the problem of false negatives[27]. In the absence of labeled data, there is a chance that the anchor cell may form a negative pair with a sample from the same class, which reduces the contribution to the contrastive loss, limiting the model’s ability to converge quickly. Moreover, in situations where samples originate from various sources or batches, the contrastive loss may erroneously treat all samples from different sources as negatives, despite them belonging to the same cell class. This failure to differentiate between domains can widen the gap between the batches[27], which could result in the inability to learn batch invariant representations, thereby hindering data integration performance.

Despite recent advances, the use of negative samples continues to pose fundamental challenges to CL. As a result, there has been an increasing interest in self-supervised learning methods that do not rely on negative samples, paving the way for further research in this area[16, 28, 29, 30]. Recently, feature decorrelation methods[30, 31, 32] were proposed in SSL to facilitate learning representations without using negative samples. The objective of these methods is to learn a decorrelated embedding space by maximizing the information content of the embeddings[30]. This prevents information collapse in the embedding variables which contain redundant information. Inspired by the efficacy of feature decorrelation based SSL in modelling unlabelled data without negative pairs[30], we expect that effective low-dimensional single cell representations can be obtained by decorrelating the different vector components of cell embeddings. However, it should be noted that using feature decorrelation-based SSL frameworks directly to discover robust representations for single cell gene profiles is sub-optimal when dealing with data from multiple sources or batches, as it fails to align them effectively. To achieve better data integration performance, we use feature decorrelation SSL coupled with unsupervised domain adaptation[33, 27]. The domain adaptation strategy is employed to bridge the domain gap between batches with varying probability distributions, improving the alignment of batches.

Here, we adopt these ideas and propose a novel *Feature Decorrelation based Representation Learning with Domain Adaptation* framework for integrative single-cell analysis. The key contributions of this paper are: 1. Learn robust cell representations of unlabelled single cell experiments in a negative-sample free self-supervised fashion, by leveraging the idea of feature decorrelation 2. Model the data integration problem using domain adaptation 3. Extend feature decorrelation based SSL to an unsupervised domain adaptation context, where batch-invariant representations are learned from unannotated samples belonging to multiple domains (or batches), possessing different probability distributions

## 2 Methods

### 2.1 Overview

scDecorr employs self-supervised feature decorrelation based representation learning coupled with domain adaptation (DA) to integrate multiple unlabelled single cell experiments. scDecorr learns cell representations in a self-supervised fashion via a joint embedding of distorted gene profiles of a cell. It accomplishes this by optimizing an objective function that maximizes similarity among the distorted embeddings while also decorrelating their components. scDecorr learns batch-invariant representations using the domain adaptation (DA) framework. It is responsible for projecting samples from multiple domains to a common manifold such that similar cell samples from all the domains lie close to each other. The DA framework learns domain(or, batch)-invariant representations by learning cell representations independently across domains by randomly sampling instances from all the domains. Due to the variations in statistical properties of samples from distinct domains, a domain-specific batch normalization (DSBN)[33] layer is used which enables better normalization and reduces the impact of domain shift on the learned representations.

### 2.2 Description

#### Representation Learning

scDecorr learns representations by using a deep network *f* _*θ*_ that projects samples from the original feature space (X) to embedding space (*Z*_*θ*_) via a joint embedding of distorted samples (Y). The network is trained with an objective such that the representations preserve as much information as possible from the presented samples while also achieving a high degree of invariance to the distortions applied[30]. This is achieved by maximising the mutual information (MI) *I*(*Z*; *X*) between the representations (Z) and presented samples (X), while minimising the MI *I*(*Z*; *Y*) between the representations (Z) and the distorted samples (Y). This can be briefly described using an objective function *L*(*θ*) := *I*(*Z*; *Y*) − *I*(*Z*; *X*). The loss can be simplified and re-written as follows *L*(*θ*) := *H*(*Z*_*θ*_|*X*) − λ*H*(*Z*_*θ*_), where H(.) denotes the entropy and λ is a positive constant. This simplified objective reveals that, the conditional entropy *Z*_*θ*_|*X* term needs to be minimized that is, *Z*_*θ*_ has to be completely determined from *X*. In other words, the information *Z*_*θ*_ contains about the distortions has to be minimized therefore, minimizing the conditional entropy loss term is equivalent to maximizing the alignment (or, similarity) between the representations of the distorted samples *Y*. As this loss term is essentially responsible for making the representations invariant to distortions applied, it is also called the *invariance term*. Furthermore, minimizing the overall objective results in maximizing the second loss term *H*(*Z*_*θ*_). Maximizing the entropy is aimed at enhancing the variability of the learned representations *Z*_*θ*_. This can be realized by minimizing the redundancy of information encoded by different vector components. To this end, the correlation between every pair of representation components are reduced towards zero, which is commonly referred to as decorrelation. The rationale behind decorrelation is to avert the occurrence of an informational collapse, wherein components may become highly correlated or vary together. The second loss term is formally designated as the *decorrelation term*.

scDecorr leverages the joint embeddings of two distorted profiles of each cell (*Z, Z*′) to calculate the invariance and covariance loss terms. Specifically, the cross-correlation matrix (W) between the embeddings is computed over a mini-batch as follows:

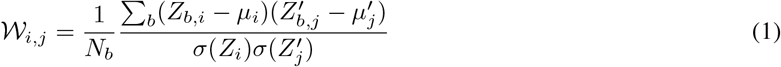

where,

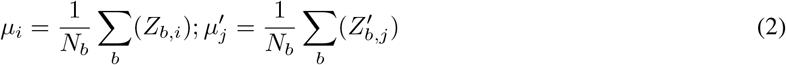

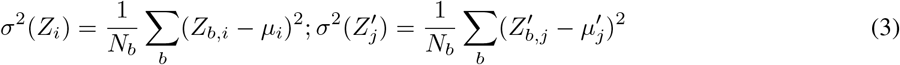

and, *b* indicates the mini-batch index and *N*_*b*_ indicates the size of the mini-batch. 𝒲_*i,j*_ values are comprised between 1 (perfect correlation) to −1 (anti-correlation), and 0 indictating uncorrelatedness.

The *invariance term* of the loss is computed using the on-diagonal elements of 𝒲, encouraging them to be close to 1

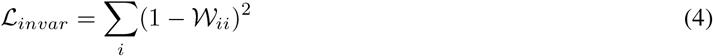

whereas the *decorrelation term* of the loss is calculated using the off-diagonal elements, encouraging them to be close to 0

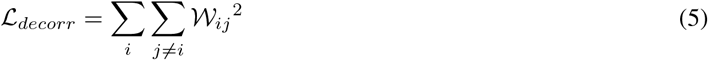

The overall loss is therefore,

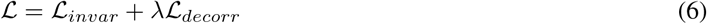

where λ is a trade-off factor balancing the invariance and decorrelation terms.

#### Domain Adaptation

However, when a mini-batch comprises of cell samples from different domains, the usual computation of the loss terms results in erroneous results. This issue arises because 𝒲, *μ*, and σ are calculated over the mini-batch. Since cell samples from different domains follow distinct probability distributions, they possess varying mean and variance values. Therefore, computing a single mean *μ* and variance σ for all the domains hurts the domain alignment performance. To address the challenge of domain differences in a mini-batch, scDecorr adopts a strategy where separate values of 𝒲^(*d*)^, *μ*^(*d*)^, and *σ*^(*d*)^ are calculated independently for each domain *d*. This is achieved through random sampling of cells from each domain, and normalizing the data independently across domain via domain-specific batch normalization layers[33]. Therefore, the loss term in Eq 6 ℒ^(*d*)^ is computed independently for every domain and the overall domain adapted loss *L*_*DA*_ is simply the mean of all the domain losses.

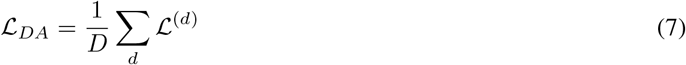

### 2.3 Method implementation for single-cell transcriptomics data

#### 2.3.1 Preprocessing

First, we remove genes expressed in less than 3 cells. Next, we log-normalize the raw gene expression counts data which involves dividing the total count by 10, 000 followed by log transformation to obtain the normalized expression matrix. Finally, we select the top 2000 highly variable genes by batch for training scDecorr. We used the Scanpy package [34] for the pre-processing steps.

#### 2.3.2 scDecorr Workflow

scDecorr accepts *D* preprocessed gene expression count matrices 𝒳 = {𝒳^(1)^, …, 𝒳^(*D*)^} from *D* distinct domains (or, sources, or batches) as input. scDecorr aims to integrate these *D* unintegrated count matrices to a common domain-invariant low-dimensional representation space Ƶ. In our experiments, the input space has a size of 2000 corresponding to the highly variable genes and the representation space has a size of 64.

##### Mini-Batch Construction

First, the mini-batch is constructed by randomly sampling *K* cells from each domain *d*. {*X*^(*d*)^ ⊆ 𝒳^(*d*)^*s.t*. ‖*X*^(*d*)^ ‖ = *K*} constitute the mini-batch specific to the domain.

##### Data Augmentation

Next, for each domain specific mini-batch *X*^(*d*)^, mini-batches of two distorted views *X*^*′*(*d*)^ and *X*^*′′*(*d*)^ are created by sampling two transformations from a distribution of data augmentations. The transformations include randomly zeroing out 20% of genes, and randomly shuffling 10% of the gene expression values.

##### Model Architecture

scDecorr has a symmetric Siamese net architecture— network consists of 2 identical branches, each responsible for embedding a particular distorted view. Each branch consists of an encoder and projector network, and their weights are shared between the branches. For every domain, the mini-batches of distorted views *X*^*′*(*d*)^ and *X*^*′′*(*d*)^ are fed to the respective branches of scDecorr. The encoder network *f*_*θ*_ produces batches of low-dimensional representations *Z*^*′*(*d*)^ = *f*_*θ*_ (*X*^*′*(*d*)^) and *Z*^*′′*(*d*)^ = *f*_*θ*_ (*X*^*′′*(*d*)^). The representations are then fed to a projector network *g*_*ϕ*_ which projects the low-dimensional representations to a high dimensional embedding space, *Y*^*‘*(*d*)^ = *g*_*ϕ*_(*Z*^*′*(*d*)^) and *Y*^*‘‘*(*d*)^ = *g*_*ϕ*_ (*Z*^*′′*(*d*)^). The output of the encoder *Z* is called *representations* and the output of the projector *Y* is called *embeddings*. The encoder consists of a densely connected (DenseNet) neural network[35, 19]. The projector consists of linear layers, the first two being followed by a ReLU and a domain-specific batch normalization layer.

##### Loss Computation

Finally, the batches of embeddings of every domain *Y*^*‘*(*d*)^ and *Y*^*‘‘*(*d*)^ are z-score normalized (along the batch axis) independently across domains using another domain-specific batch-normalization layer with no affine transform. This ensures that every embedding component of a domain has a mean of 0 and variance of 1 over its mini-batch so that the domain specific cross-correlation matrix can be computed simply by a matrix multiplication between *Y*^*‘*(*d*)^ and *Y*^*‘‘*(*d*)^:

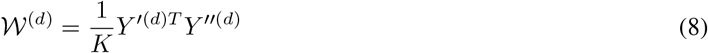

The domain loss ℒ^(*d*)^ is then computed from 𝒲^(*d*)^ as follows:

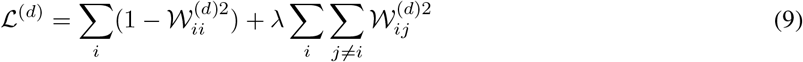

The overall loss can simply be computed as the average of all the domain losses.

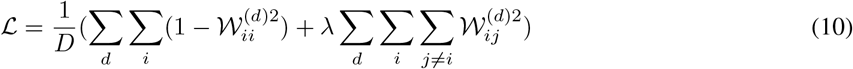

##### Model Optimization

We train scDecorr for 1000 epochs using the Adam optimizer with a base learning rate of 0.0003. Cosine annealing with a warmup of 10 epochs is used to anneal the learning rate over time. Early stopping is also used to stop training if there is no improvement in loss for 50 epochs.

After training scDecorr on the unintegrated dataset 𝒳, the integrated 64 dimensional representations Ƶ = *f* (𝒳) are used as features for downstream tasks.

#### 2.3.3 Downstream Tasks

We use scDecorr for three downstream tasks namely clustering, batch correction (batch-mixing), and reference to query label annotation.

##### Clustering

We cluster the feature representations using the Leiden community detection algorithm with a resolution of 0.2. We aim to detect clusters which group functionally similar cells together.

##### Batch Mixing

The presence of batch effects becomes a significant confounding factor in cell-type clustering when integrating multiple scRNA-seq datasets originating from different sequencing batches or technologies. To address this, we first integrate the datasets to obtain a batch-invariant representation and then feed it to the clustering algorithm.

##### Label Annotation

For annotating an unlabelled query dataset, we first integrate the reference and query datasets together using scDecorr. Next, a XGBoost model is trained on the features extracted from the reference. The trained XGBoost model is then used to predict cell types of the query dataset. However, it is to be noted that, the cells of the query dataset having cell types not present in the reference dataset can not be annotated by this method.

## 3 Results

### 3.1 Dataset Description

We benchmark scDecorr on 4 real world benchmark scRNA-seq datasets. These datasets encompass a range of data integration tasks such as integration across donors, chemistries, platforms, and integration of two or more batches. They also encompass diverse tissues, cell types and species, such as human immune cells, human kidney cells, and mouse cells. The cell type annotations and batch labels of these datasets are pre-existing and already available from original publications. The cell-type annotations are not required for training scDecorr, but are used in the downstream tasks. The dataset details are shown in Table 1.

**Table 1:**
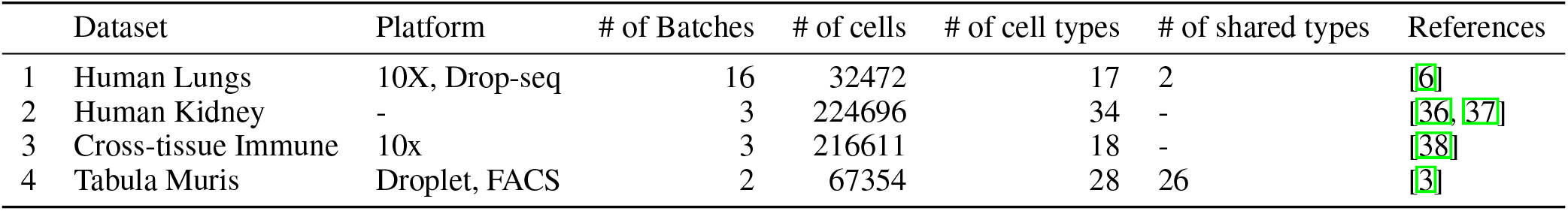
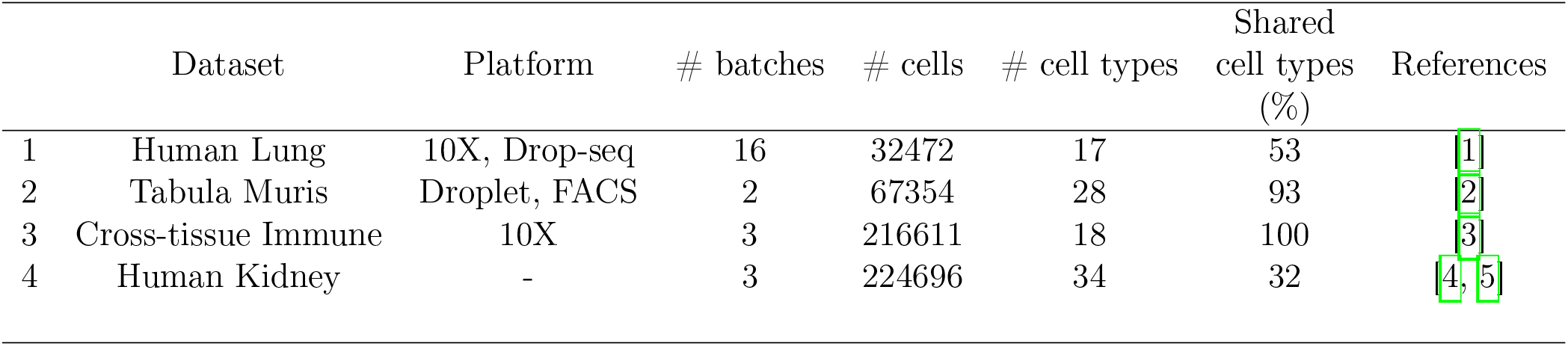
Datasets details - overview of datasets used in this study.

### 3.2 scDecorr Training Configuration

In our experiments, we use two different configurations *S* and *L* for training scDecorr. Configuration S is used for small datasets (with circa 50k cells or less) and configuration L for large-scale datasets (with circa 100k cells or more). For configuration *S*, a mini-batch size of *K* = 512 is used. An 11 layer DenseNet model and a 512 − 512 − 512 layer MLP are used as the encoder and decoder respectively. Furthermore, for configuration *L*, a mini-batch size of *K* = 2048 is used. A 21 layer DenseNet model[35] and a 1024 − 1024 − 1024 layer MLP are used as the encoder and decoder respectively. The best training configurations of scDecorr for all the benchmark datasets are shown in Table2.

**Table 2:**
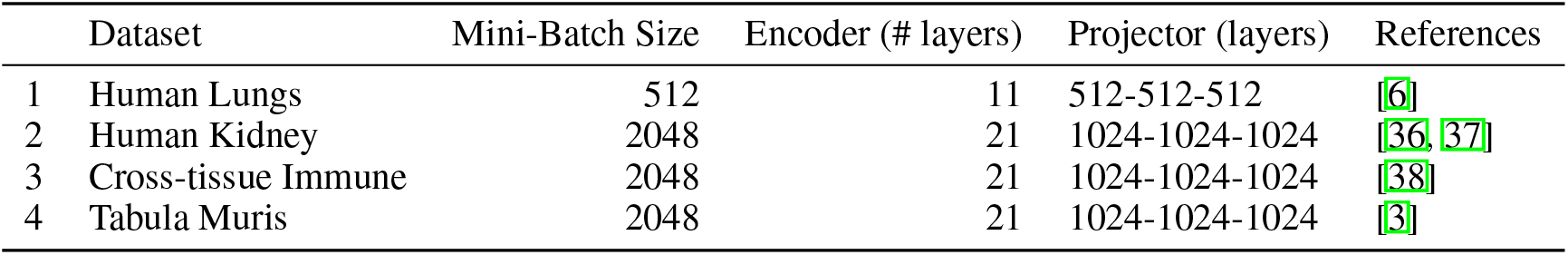
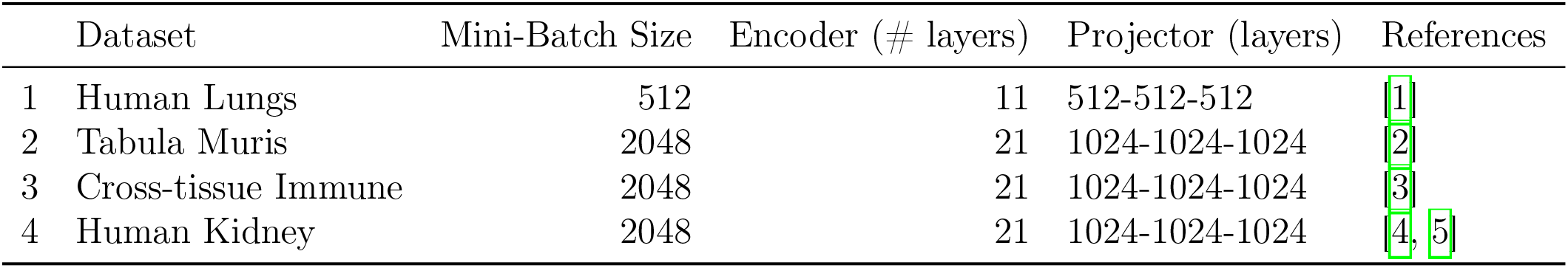
Best training configurations of scDecorr for every dataset.

**Table 3:** Human Lungs: Benchmark Results of Data Integration across Donors.

|  | PCA | LIGER | Scanorama | Harmony | scVI | scDecorr |
| --- | --- | --- | --- | --- | --- | --- |
| Metrics |  |  |  |  |  |  |
| leiden.ARI | 0.489 | 0.357 | 0.107 | 0.497 | 0.541 | 0.636 |
| leiden.NMI | 0.735 | 0.593 | 0.326 | 0.659 | 0.728 | 0.726 |
| silhouette | 0.554 | 0.531 | 0.433 | 0.535 | 0.543 | 0.623 |
| S_bio | 0.593 | 0.494 | 0.289 | 0.564 | 0.604 | 0.662 |
| batch_asw | 0.882 | 0.854 | 0.854 | 0.894 | 0.902 | 0.810 |
| kbet | 0.230 | 0.535 | 0.298 | 0.460 | 0.342 | 0.339 |
| S_batch | 0.556 | 0.694 | 0.576 | 0.677 | 0.622 | 0.574 |
| S_overall | 0.582 | 0.554 | 0.375 | 0.598 | 0.609 | 0.635 |

**Table 4:**
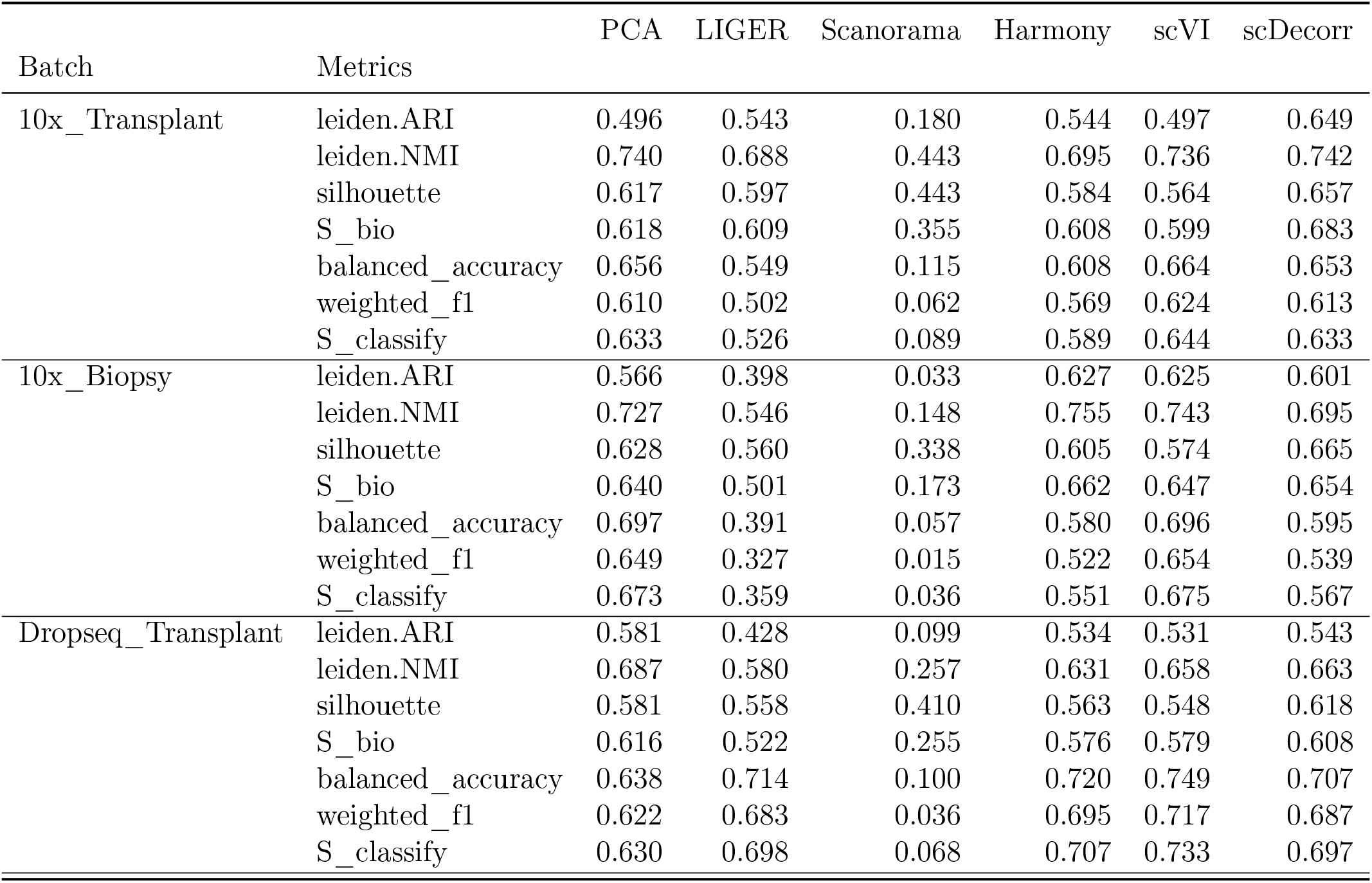
Human Lungs: Experiment-wise Label Transferring Results.

| Batch | Metrics | PCA | LIGER | Scanorama | Harmony | scVI | scDecorr |
| --- | --- | --- | --- | --- | --- | --- | --- |
| 10x_Transplant | leiden.ARI | 0.496 | 0.543 | 0.180 | 0.544 | 0.497 | 0.649 |
|  | leiden.NMI | 0.740 | 0.688 | 0.443 | 0.695 | 0.736 | 0.742 |
|  | silhouette | 0.617 | 0.597 | 0.443 | 0.584 | 0.564 | 0.657 |
|  | S_bio | 0.618 | 0.609 | 0.355 | 0.608 | 0.599 | 0.683 |
|  | balanced_accuracy | 0.656 | 0.549 | 0.115 | 0.608 | 0.664 | 0.653 |
|  | weighted_f1 | 0.610 | 0.502 | 0.062 | 0.569 | 0.624 | 0.613 |
|  | S_classify | 0.633 | 0.526 | 0.089 | 0.589 | 0.644 | 0.633 |
| 10x_Biopsy | leiden.ARI | 0.566 | 0.398 | 0.033 | 0.627 | 0.625 | 0.601 |
|  | leiden.NMI | 0.727 | 0.546 | 0.148 | 0.755 | 0.743 | 0.695 |
|  | silhouette | 0.628 | 0.560 | 0.338 | 0.605 | 0.574 | 0.665 |
|  | S_bio | 0.640 | 0.501 | 0.173 | 0.662 | 0.647 | 0.654 |
|  | balanced_accuracy | 0.697 | 0.391 | 0.057 | 0.580 | 0.696 | 0.595 |
|  | weighted_f1 | 0.649 | 0.327 | 0.015 | 0.522 | 0.654 | 0.539 |
|  | S_classify | 0.673 | 0.359 | 0.036 | 0.551 | 0.675 | 0.567 |
| Dropseq_Transplant | leiden.ARI | 0.581 | 0.428 | 0.099 | 0.534 | 0.531 | 0.543 |
|  | leiden.NMI | 0.687 | 0.580 | 0.257 | 0.631 | 0.658 | 0.663 |
|  | silhouette | 0.581 | 0.558 | 0.410 | 0.563 | 0.548 | 0.618 |
|  | S_bio | 0.616 | 0.522 | 0.255 | 0.576 | 0.579 | 0.608 |
|  | balanced_accuracy | 0.638 | 0.714 | 0.100 | 0.720 | 0.749 | 0.707 |
|  | weighted_f1 | 0.622 | 0.683 | 0.036 | 0.695 | 0.717 | 0.687 |
|  | S_classify | 0.630 | 0.698 | 0.068 | 0.707 | 0.733 | 0.697 |

**Table 5:** Kidney: Benchmark Results of Data Integration across Datasets.

|  | PCA | Scanorama | LIGER | Harmony | trVAE | scVI | scDecorr |
| --- | --- | --- | --- | --- | --- | --- | --- |
| Metrics |  |  |  |  |  |  |  |
| leiden.ARI | 0.681 | 0.021 | 0.406 | 0.452 | 0.524 | 0.563 | 0.654 |
| leiden.NMI | 0.778 | 0.062 | 0.586 | 0.615 | 0.652 | 0.696 | 0.755 |
| silhouette | 0.561 | 0.457 | 0.455 | 0.492 | 0.504 | 0.525 | 0.579 |
| S_bio | 0.673 | 0.180 | 0.483 | 0.520 | 0.560 | 0.595 | 0.663 |
| batch_asw | 0.853 | 0.839 | 0.905 | 0.942 | 0.872 | 0.929 | 0.851 |
| wisi | 0.102 | 0.221 | 0.360 | 0.314 | 0.176 | 0.191 | 0.128 |
| entropy | 0.100 | 0.227 | 0.373 | 0.307 | 0.178 | 0.187 | 0.129 |
| S_batch | 0.352 | 0.429 | 0.546 | 0.521 | 0.409 | 0.436 | 0.369 |
| S_overall | 0.577 | 0.255 | 0.502 | 0.520 | 0.515 | 0.547 | 0.575 |

**Table 6:**
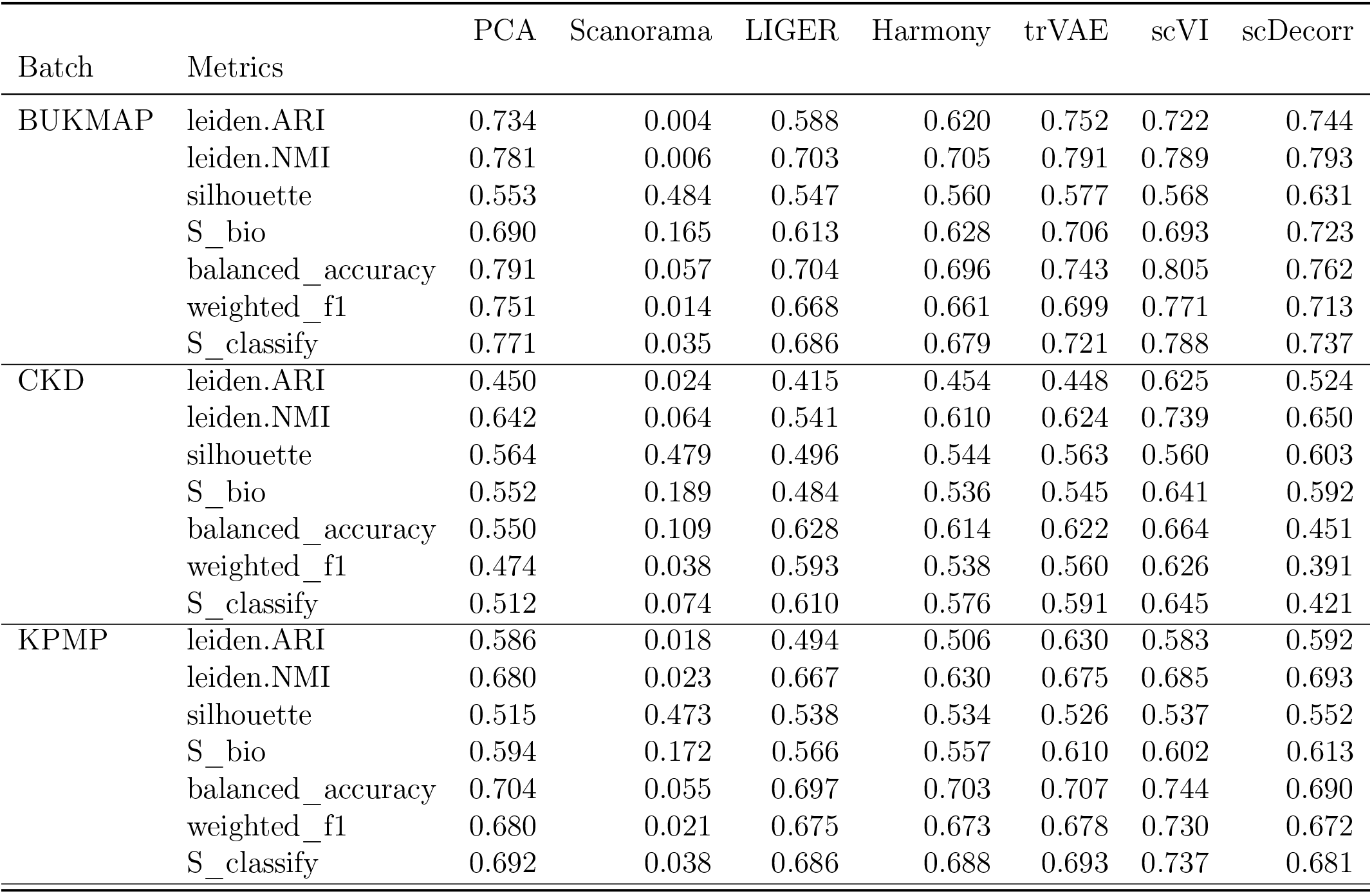
Kidney: Label Transferring Results across Datasets.

|  |  | PCA | Scanorama | LIGER | Harmony | trVAE | scVI | scDecorr |
| --- | --- | --- | --- | --- | --- | --- | --- | --- |
| Batch | Metrics |  |  |  |  |  |  |  |
| BUKMAP | leiden.ARI | 0.734 | 0.004 | 0.588 | 0.620 | 0.752 | 0.722 | 0.744 |
|  | leiden.NMI | 0.781 | 0.006 | 0.703 | 0.705 | 0.791 | 0.789 | 0.793 |
|  | silhouette | 0.553 | 0.484 | 0.547 | 0.560 | 0.577 | 0.568 | 0.631 |
|  | S_bio | 0.690 | 0.165 | 0.613 | 0.628 | 0.706 | 0.693 | 0.723 |
|  | balanced_accuracy | 0.791 | 0.057 | 0.704 | 0.696 | 0.743 | 0.805 | 0.762 |
|  | weighted_f1 | 0.751 | 0.014 | 0.668 | 0.661 | 0.699 | 0.771 | 0.713 |
|  | S_classify | 0.771 | 0.035 | 0.686 | 0.679 | 0.721 | 0.788 | 0.737 |
| CKD | leiden.ARI | 0.450 | 0.024 | 0.415 | 0.454 | 0.448 | 0.625 | 0.524 |
|  | leiden.NMI | 0.642 | 0.064 | 0.541 | 0.610 | 0.624 | 0.739 | 0.650 |
|  | silhouette | 0.564 | 0.479 | 0.496 | 0.544 | 0.563 | 0.560 | 0.603 |
|  | S_bio | 0.552 | 0.189 | 0.484 | 0.536 | 0.545 | 0.641 | 0.592 |
|  | balanced_accuracy | 0.550 | 0.109 | 0.628 | 0.614 | 0.622 | 0.664 | 0.451 |
|  | weighted_f1 | 0.474 | 0.038 | 0.593 | 0.538 | 0.560 | 0.626 | 0.391 |
|  | S_classify | 0.512 | 0.074 | 0.610 | 0.576 | 0.591 | 0.645 | 0.421 |
| KPMP | leiden.ARI | 0.586 | 0.018 | 0.494 | 0.506 | 0.630 | 0.583 | 0.592 |
|  | leiden.NMI | 0.680 | 0.023 | 0.667 | 0.630 | 0.675 | 0.685 | 0.693 |
|  | silhouette | 0.515 | 0.473 | 0.538 | 0.534 | 0.526 | 0.537 | 0.552 |
|  | S_bio | 0.594 | 0.172 | 0.566 | 0.557 | 0.610 | 0.602 | 0.613 |
|  | balanced_accuracy | 0.704 | 0.055 | 0.697 | 0.703 | 0.707 | 0.744 | 0.690 |
|  | weighted_f1 | 0.680 | 0.021 | 0.675 | 0.673 | 0.678 | 0.730 | 0.672 |
|  | S_classify | 0.692 | 0.038 | 0.686 | 0.688 | 0.693 | 0.737 | 0.681 |

**Table 7:** Crosstissue-Immune: Benchmark Results of Data Integration across Chemistries.

|  | PCA | Scanorama | LIGER | Harmony | trVAE | scVI | scDecorr |
| --- | --- | --- | --- | --- | --- | --- | --- |
| Metrics |  |  |  |  |  |  |  |
| leiden.ARI | 0.201 | 0.001 | 0.287 | 0.323 | 0.293 | 0.276 | 0.423 |
| leiden.NMI | 0.407 | 0.003 | 0.438 | 0.528 | 0.525 | 0.462 | 0.645 |
| silhouette | 0.516 | 0.493 | 0.514 | 0.517 | 0.519 | 0.515 | 0.559 |
| S_bio | 0.375 | 0.166 | 0.413 | 0.456 | 0.446 | 0.418 | 0.542 |
| batch_asw | 0.840 | 0.979 | 0.898 | 0.939 | 0.943 | 0.951 | 0.836 |
| wisi | 0.014 | 0.318 | 0.402 | 0.248 | 0.372 | 0.188 | 0.186 |
| entropy | 0.014 | 0.329 | 0.425 | 0.258 | 0.392 | 0.193 | 0.195 |
| S_batch | 0.289 | 0.542 | 0.575 | 0.482 | 0.569 | 0.444 | 0.406 |
| S_overall | 0.349 | 0.279 | 0.461 | 0.464 | 0.483 | 0.425 | 0.501 |

**Table 8:** Crosstissue-Immune: Label Transferring Results across Chemistries.

|  |  | PCA | Scanorama | LIGER | Harmony | trVAE | scVI | scDecorr |
| --- | --- | --- | --- | --- | --- | --- | --- | --- |
| Batch | Metrics |  |  |  |  |  |  |  |
| 5v1 | leiden.ARI | 0.209 | 0.001 | 0.239 | 0.224 | 0.170 | 0.087 | 0.375 |
|  | leiden.NMI | 0.419 | 0.006 | 0.358 | 0.437 | 0.376 | 0.208 | 0.582 |
|  | silhouette | 0.506 | 0.492 | 0.498 | 0.506 | 0.517 | 0.508 | 0.518 |
|  | S_bio | 0.378 | 0.167 | 0.365 | 0.389 | 0.355 | 0.268 | 0.492 |
|  | balanced_accuracy | 0.684 | 0.057 | 0.627 | 0.690 | 0.717 | 0.751 | 0.770 |
|  | weighted_f1 | 0.680 | 0.012 | 0.623 | 0.685 | 0.715 | 0.751 | 0.770 |
|  | S_classify | 0.682 | 0.034 | 0.625 | 0.687 | 0.716 | 0.751 | 0.770 |
| 3 | leiden.ARI | 0.377 | 0.000 | 0.367 | 0.389 | 0.388 | 0.320 | 0.484 |
|  | leiden.NMI | 0.565 | 0.002 | 0.585 | 0.577 | 0.560 | 0.523 | 0.675 |
|  | silhouette | 0.530 | 0.495 | 0.542 | 0.524 | 0.521 | 0.518 | 0.588 |
|  | S_bio | 0.491 | 0.166 | 0.498 | 0.497 | 0.489 | 0.454 | 0.582 |
|  | balanced_accuracy | 0.693 | 0.056 | 0.699 | 0.776 | 0.751 | 0.758 | 0.790 |
|  | weighted_f1 | 0.679 | 0.015 | 0.696 | 0.776 | 0.747 | 0.745 | 0.772 |
|  | S_classify | 0.686 | 0.036 | 0.697 | 0.776 | 0.749 | 0.751 | 0.781 |
| 5v2 | leiden.ARI | 0.328 | 0.000 | 0.248 | 0.312 | 0.300 | 0.239 | 0.439 |
|  | leiden.NMI | 0.471 | 0.002 | 0.405 | 0.469 | 0.481 | 0.360 | 0.632 |
|  | silhouette | 0.521 | 0.495 | 0.501 | 0.520 | 0.525 | 0.520 | 0.569 |
|  | S_bio | 0.440 | 0.166 | 0.385 | 0.434 | 0.435 | 0.373 | 0.547 |
|  | balanced_accuracy | 0.723 | 0.057 | 0.611 | 0.754 | 0.754 | 0.795 | 0.830 |
|  | weighted_f1 | 0.717 | 0.012 | 0.599 | 0.753 | 0.757 | 0.794 | 0.830 |
|  | S_classify | 0.720 | 0.035 | 0.605 | 0.753 | 0.755 | 0.795 | 0.830 |

**Table 9:** Tabula Muris: Benchmark Results of Data Integration across Experiments.

|  | PCA | Scanorama | LIGER | Harmony | trVAE | scVI | scDecorr |
| --- | --- | --- | --- | --- | --- | --- | --- |
| Metrics |  |  |  |  |  |  |  |
| leiden.ARI | 0.641 | -0.000 | 0.564 | 0.669 | 0.672 | 0.685 | 0.676 |
| leiden.NMI | 0.792 | 0.001 | 0.770 | 0.807 | 0.826 | 0.833 | 0.803 |
| silhouette | 0.582 | 0.496 | 0.600 | 0.569 | 0.571 | 0.546 | 0.591 |
| S_bio | 0.672 | 0.166 | 0.645 | 0.682 | 0.690 | 0.688 | 0.690 |
| batch_asw | 0.739 | 0.992 | 0.805 | 0.869 | 0.826 | 0.894 | 0.777 |
| kbet | 0.581 | 0.229 | 0.879 | 0.797 | 0.802 | 0.798 | 0.856 |
| S_batch | 0.660 | 0.611 | 0.842 | 0.833 | 0.814 | 0.846 | 0.816 |
| S_overall | 0.668 | 0.299 | 0.704 | 0.727 | 0.727 | 0.735 | 0.728 |

**Table 10:**
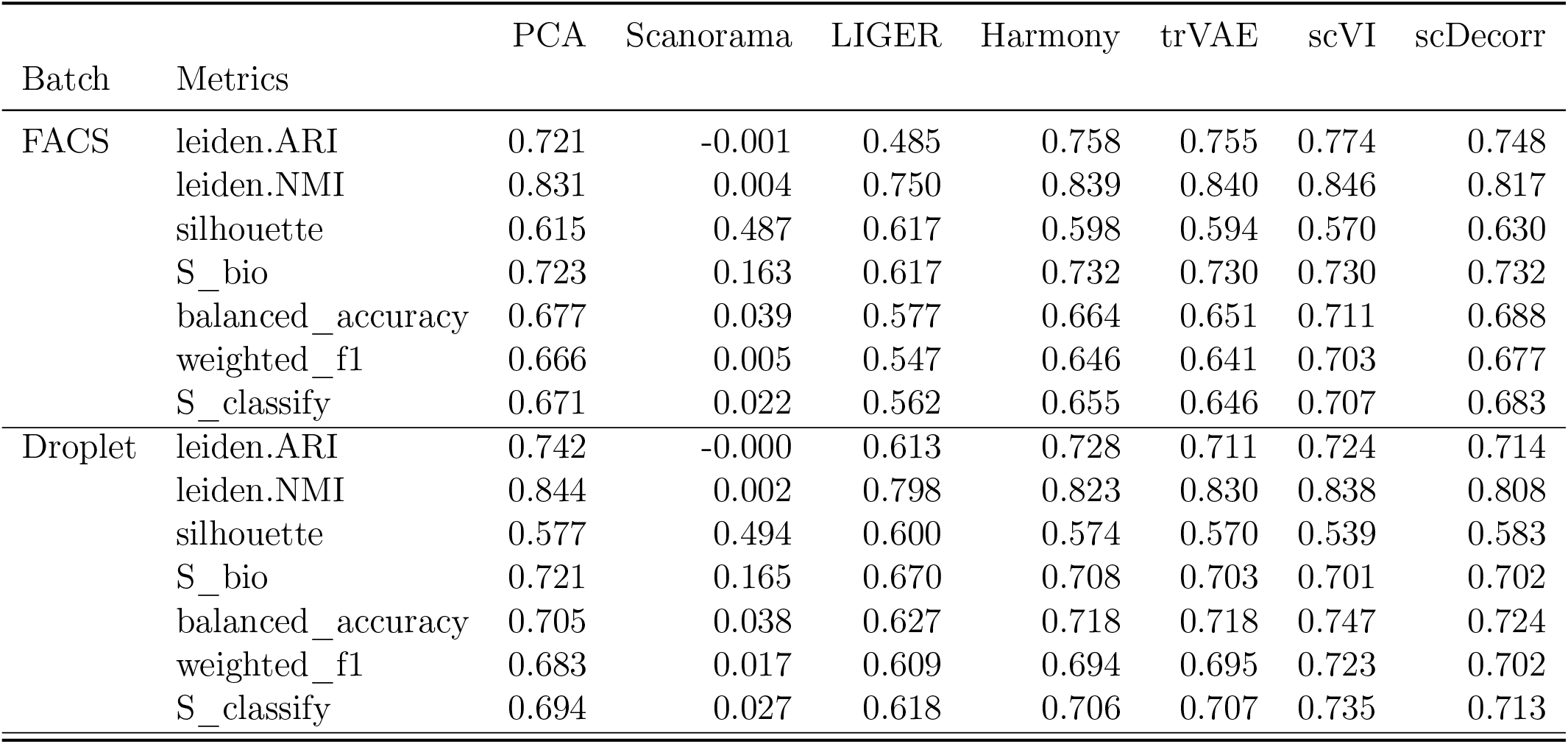
Tabula Muris: Experiment-wise Label Transferring Results.

|  |  | PCA | Scanorama | LIGER | Harmony | trVAE | scVI | scDecorr |
| --- | --- | --- | --- | --- | --- | --- | --- | --- |
| Batch | Metrics |  |  |  |  |  |  |  |
| FACS | leiden.ARI | 0.721 | -0.001 | 0.485 | 0.758 | 0.755 | 0.774 | 0.748 |
|  | leiden.NMI | 0.831 | 0.004 | 0.750 | 0.839 | 0.840 | 0.846 | 0.817 |
|  | silhouette | 0.615 | 0.487 | 0.617 | 0.598 | 0.594 | 0.570 | 0.630 |
|  | S_bio | 0.723 | 0.163 | 0.617 | 0.732 | 0.730 | 0.730 | 0.732 |
|  | balanced_accuracy | 0.677 | 0.039 | 0.577 | 0.664 | 0.651 | 0.711 | 0.688 |
|  | weighted_f1 | 0.666 | 0.005 | 0.547 | 0.646 | 0.641 | 0.703 | 0.677 |
|  | S_classify | 0.671 | 0.022 | 0.562 | 0.655 | 0.646 | 0.707 | 0.683 |
| Droplet | leiden.ARI | 0.742 | -0.000 | 0.613 | 0.728 | 0.711 | 0.724 | 0.714 |
|  | leiden.NMI | 0.844 | 0.002 | 0.798 | 0.823 | 0.830 | 0.838 | 0.808 |
|  | silhouette | 0.577 | 0.494 | 0.600 | 0.574 | 0.570 | 0.539 | 0.583 |
|  | S_bio | 0.721 | 0.165 | 0.670 | 0.708 | 0.703 | 0.701 | 0.702 |
|  | balanced_accuracy | 0.705 | 0.038 | 0.627 | 0.718 | 0.718 | 0.747 | 0.724 |
|  | weighted_f1 | 0.683 | 0.017 | 0.609 | 0.694 | 0.695 | 0.723 | 0.702 |
|  | S_classify | 0.694 | 0.027 | 0.618 | 0.706 | 0.707 | 0.735 | 0.713 |

### 3.3 Experimental Setup

To evaluate the performance of scDecorr, we assess its effectiveness in two different downstream scenarios: data integration and label transfer. To establish a benchmark, we compare scDecorr with six other methods: Harmony[10], scVI[12], trVAE[39], LIGER[40], Scanorama[41], and PCA[42]. For our experiments, we utilize the default implementations of these methods available in the scib package [6]. For generating LIGER representations, we employ the default implementation from the pyLIGER package [43]. Subsequently, the batch-integrated feature representations extracted by these methods serve as the input for the downstream tasks.

#### 3.3.1 Data Integration

This experiment aims to evaluate the performance of the method in preserving biological signals and correcting batch effects. The evaluation metrics are computed using the scib package[6], unless otherwise specified.

**Clustering (Biological conservation)** is assessed by measuring cell-type clustering on the batch-integrated feature representations obtained from each method. Three metrics, namely Adjusted Rand Index (ARI)[44], Normalized Mutual Information (NMI)[45, 6], and cell-type silhouette (cSil) score[46, 6], are utilized to evaluate the clustering performance.

The Adjusted Rand Index (ARI)[44] quantifies the extent of agreement between the clustering labels and the ground truth cell-type labels, considering the agreement beyond what would be expected by chance. A higher ARI value, ranging from −1 to 1, indicates a better agreement between the predicted clusters and the true labels, with 1 indicating a perfect match.

The Normalized Mutual Information (NMI)[45, 6] measures the mutual information between the predicted clusters and the true labels, normalized by the entropy of the predicted and true label distributions. NMI values range from 0 to 1, with 1 indicating a perfect match between the predicted clusters and the true labels.

The cell-type silhouette score (cSil)[46, 6] evaluates the degree of separation between the clusters. It assesses how well each cell is assigned to its corresponding cluster compared to other clusters. The silhouette score ranges from −1 to 1, with higher values indicating better separation between clusters and overall improved clustering performance. A score close to 1 implies well-separated and distinct clusters, while a score close to −1 suggests overlapping or poorly separated clusters.

To enable easy comparison among the benchmark methods, the ARI, NMI, and cSil scores are scaled to a range of 0 to 1. The overall biological conservation performance *S*_*bio*_ of a method on a scRNA-seq dataset is calculated by taking the simple mean of these three scores.

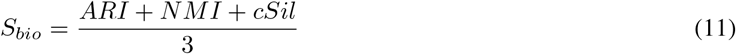

**Batch-effect correction** is assessed by computing several scores, including batch ASW (bASW)[47], kBET[47], weighted Gini-Simpson index (wGSI), and Shannon-Wiener entropy score, on the integrated feature representations.

The average silhouette width with batch labels (bASW)[47] measures batch cluster mixing by determining the silhouette width for each cell type using batch labels and then averaging across all cell types. A value of 1 indicates an ideal mixing scenario, while 0 suggests strong separation between batches.

kBET[47] evaluates batch mixing by comparing the local batch label distribution among randomly selected nearest-neighbor cells with the global batch label distribution. It is scaled to a range of 0 to 1 for easier comparison.

The Shannon-Wiener entropy[48, 5] score quantifies the diversity in the distribution of cells across different batches. The entropy value can range from 0 (indicating a completely uniform distribution) to a maximum value determined by the number of batches and the distribution of cells among them. To scale the entropy score to a range of 0 to 1, it is divided by the logarithm of the number of batches.

The Gini-Simpson index[49, 50], commonly used in ecology and biodiversity studies, is adapted to evaluate batch mixing. It is the complement of Simpson’s index and quantifies the probability that two randomly selected entities from a population belong to different categories (e.g., batches). The index ranges from 0 to 1, where 0 indicates minimum diversity and 1 indicates maximum diversity. In this evaluation, the Gini-Simpson index incorporates a weighting term that considers the nearest neighbor distances between cells from the same batch.

For smaller datasets (≤70*k* cells), namely Human Lung and Tabula Muris, bASW and kBET are used to evaluate batch mixing performance. However, for larger datasets (≥: 200*k* cells), namely Human Kidney and Crosstissue Immune, the scib implementation of kBET encountered memory issues. Therefore, entropy and weighted Gini-Simpson index (wGSI) are utilized for these datasets instead of kBET.

The overall batch-correction performance *S*_*batch*_ of the benchmark methods is calculated using the following equations:

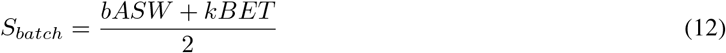

for Datasets 1, and 4,

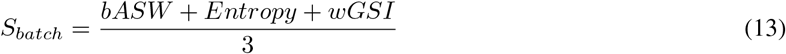

for Datasets 2, 3. Since the scib package does not provide implementations of entropy and wGSI scores, we independently implement them.

Following [6, 5], the overall data integration performance of a benchmark method on a scRNA-seq dataset is evaluated using,

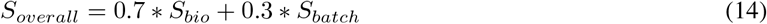

The aggregation for integration score assigns a higher weight to the clustering (biological conservation) scores, highlighting the greater significance of preserving biological information compared to batch mixing.

#### 3.3.2 Label Transferring

The process of label transferring involves assigning cell-type labels to an unlabeled query dataset using information from a reference dataset. In the case of scRNA-seq data, where gene expression data originates from multiple sources such as different platforms, datasets, or batches, we evaluate the performance of benchmark methods in annotating cell types across different sources. To accomplish this, we treat cells from a particular source as the query dataset, while considering the rest of the cells as the reference dataset.

Assuming that the data has already been integrated using a specific method, we evaluate the label annotation performance through the following approaches:

##### Clustering (Biological Conservation) of Query

Initially, we apply the Leiden clustering algorithm to the batch-integrated representations of the entire dataset. This enables us to evaluate the biological conservation quality of the query cells. We compute metric scores such as ARI (Adjusted Rand Index), NMI (Normalized Mutual Information), and cSil (cell-type silhouette) by comparing the cluster labels assigned by Leiden clustering to the ground truth cell-type labels of the query cells. This evaluation strategy is particularly valuable for datasets with minimal shared cell types between the reference and query datasets. The overall clustering performance of a method on the query cells is determined using the *S*_*bio*_ score (see Equation 11).

##### Classification of Query

Next, we train an XGBoost model using the features of the reference cells and evaluate its performance on the query cells. Only the common cell types between the reference and query datasets are used for training and evaluation. The label annotation classification performance is assessed using metrics such as balanced accuracy and weighted F1 scores. This evaluation strategy is particularly useful for datasets with a substantial number of shared cell types between the reference and query datasets. The overall classification performance of a method on the query cells is computed using the *S*_*classify*_ score.

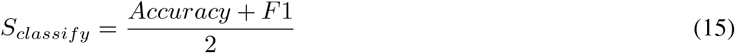

In summary, we evaluate the inter-source label annotation performance of benchmark methods by assessing their performance in clustering and classifying the query cells using reference information.

### 3.4 Visualization

To visualize the data integration process, we employ UMAP embeddings [51] of the batch-integrated feature representations, generated by a specific method. The label annotation UMAP plot of the complete dataset is constructed by aggregating the predicted cell-type labels from each query batch. These labels are obtained using a leave-one-out label transfer scheme (Refer to Label Annotation in Sec 2.3.3).

### 3.5 Use cases

In this study, we present the application of scDecorr to four different datasets: Human Lungs[6], Human Kidney[36, 37], Tabula Muris[3], and Cross-tissue Immune[38]. These datasets exhibit variations in terms of the number of cells, cell types, and batches(refer to Table 1). We assess the effectiveness of scDecorr through two distinct use cases:

- **Use Case 1**: Integrating datasets with minimal overlap in cell types across batches. This involves the Human Lung and Human Kidney datasets, which consist of 32,472 and 224,696 cells, respectively.
- **Use Case 2**: Integrating datasets with a significant proportion of shared cell types across batches. We examine this scenario using the Tabula Muris and Cross-tissue Immune datasets, comprising 67,354 and 216,611 cells, respectively. Since these two datasets include single cells sequenced from various organs, we employ them to investigate the performance of inter-organ label transfer using scDecorr.

These use cases encompass a wide range of scRNA-seq datasets that differ in scale and originate from diverse platforms, studies, donors, and technology chemistries. Furthermore, the datasets include scenarios with both two and multiple batches.

#### 3.5.1 Case Study 1: Benchmarking scDecorr on datasets with minimal shared cell types across batches Integrating Human Lung dataset across donors from three batches

The Human Lung dataset[6], comprises a comprehensive collection of 32,472 cells obtained from three distinct studies: 10x (Transplant), 10x (Biopsy), and Drop-seq (Transplant). Each study comprises of cells obtained from multiple donors. Specifically, the 10x (Transplant) study includes 12,725 single cells sequenced from six donors identified as 1, 2, …, 6. The 10x (Biopsy) study encompasses 10,046 cells sequenced from six donors identified as *A*1, *A*2, …, *A*6. Lastly, the Drop-seq (Transplant) study consists of 9,701 cells originating from four donors identified as *B*1, *B*2, …, *B*4. These 16 unique donor identifiers represent the various batches within this dataset, resulting in a total of 16 batches. The cell-type distribution within the 10x-Biopsy study (donors A1-A6) exhibits significant variation compared to the other studies. This specific group predominantly consists of Ciliated, Basal-1, Basal-2, and Secretory cell types, which either appear in minor proportions or are entirely absent in the remaining batches (refer to Figure 3).

**Figure 1.**
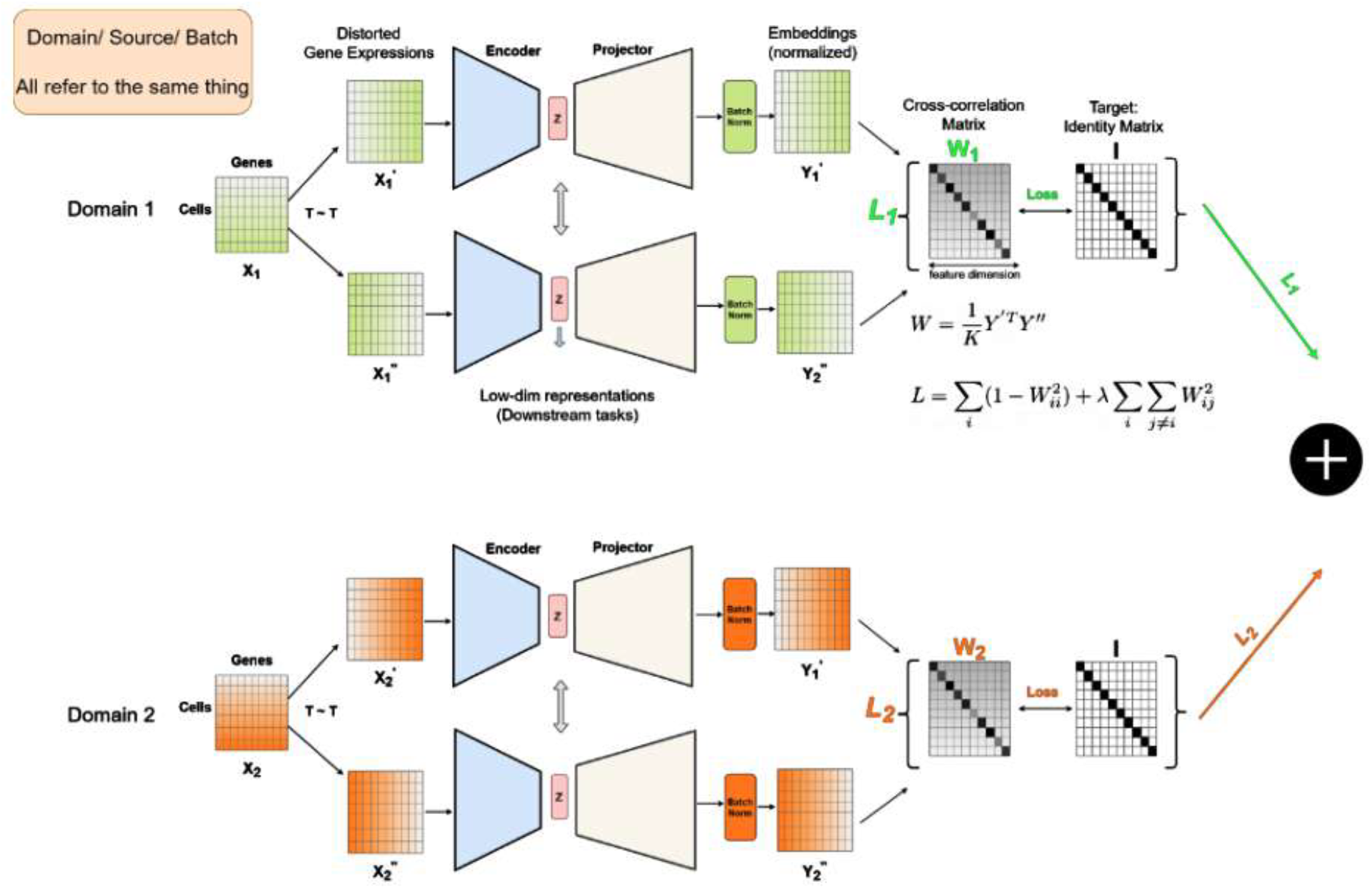
Overview of the scDecorr workflow - scDecorr takes as input single-cell gene-expression matrix coming from different studies (Domains) and uses a self-supervised feature decorrelation approach using a siamese twin model to obtain an optimal data representation. This representation is then used to perform typical single-cell downstream tasks such as clustering, batch-effect correction and cell-type annotation. We show the utility of scDecorr on 4 datasets from different tissues and techonolgoies and compare it to other computational tools on similar tasks.

**Figure 2.**
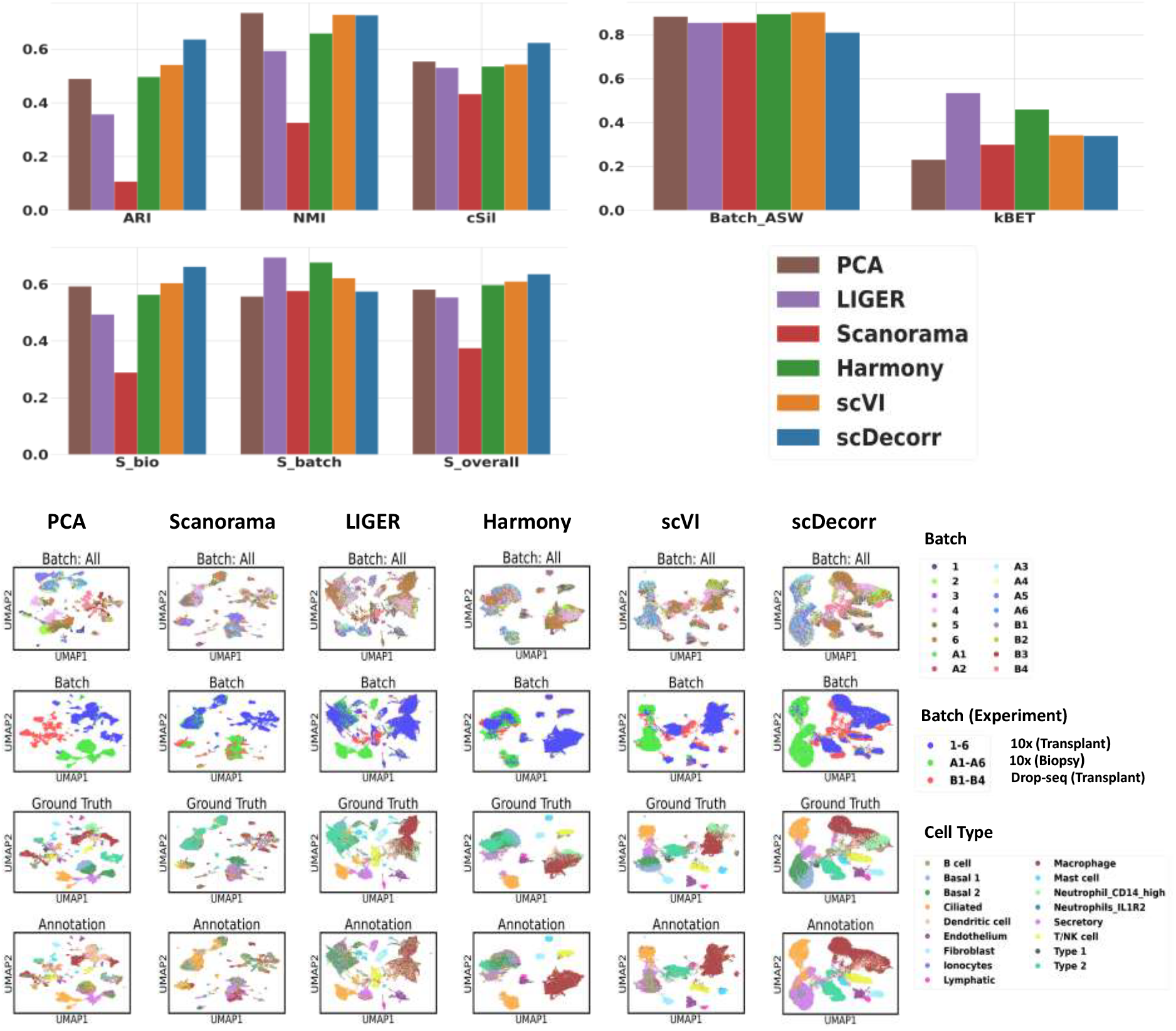
Human Lung data - Integration Benchmark Results. The lung dataset consists of 16 samples from 10X Transplant (n=6) and biopsy samples (n=6), and Drop-seq Transplant (n=4) experiments. Ground Truth refers to author’s annotation given in the original publication. scDecorr has the highest *S*_*bio*_ and overall *S*_*overall*_ score (Table S3). When compared to clustering given in the original publication, scDecorr has an ARI (Adjusted Rand Index) and cell-type silhouette score of 0.636 and 0.623, respectively (Table S3).

**Figure 3.**
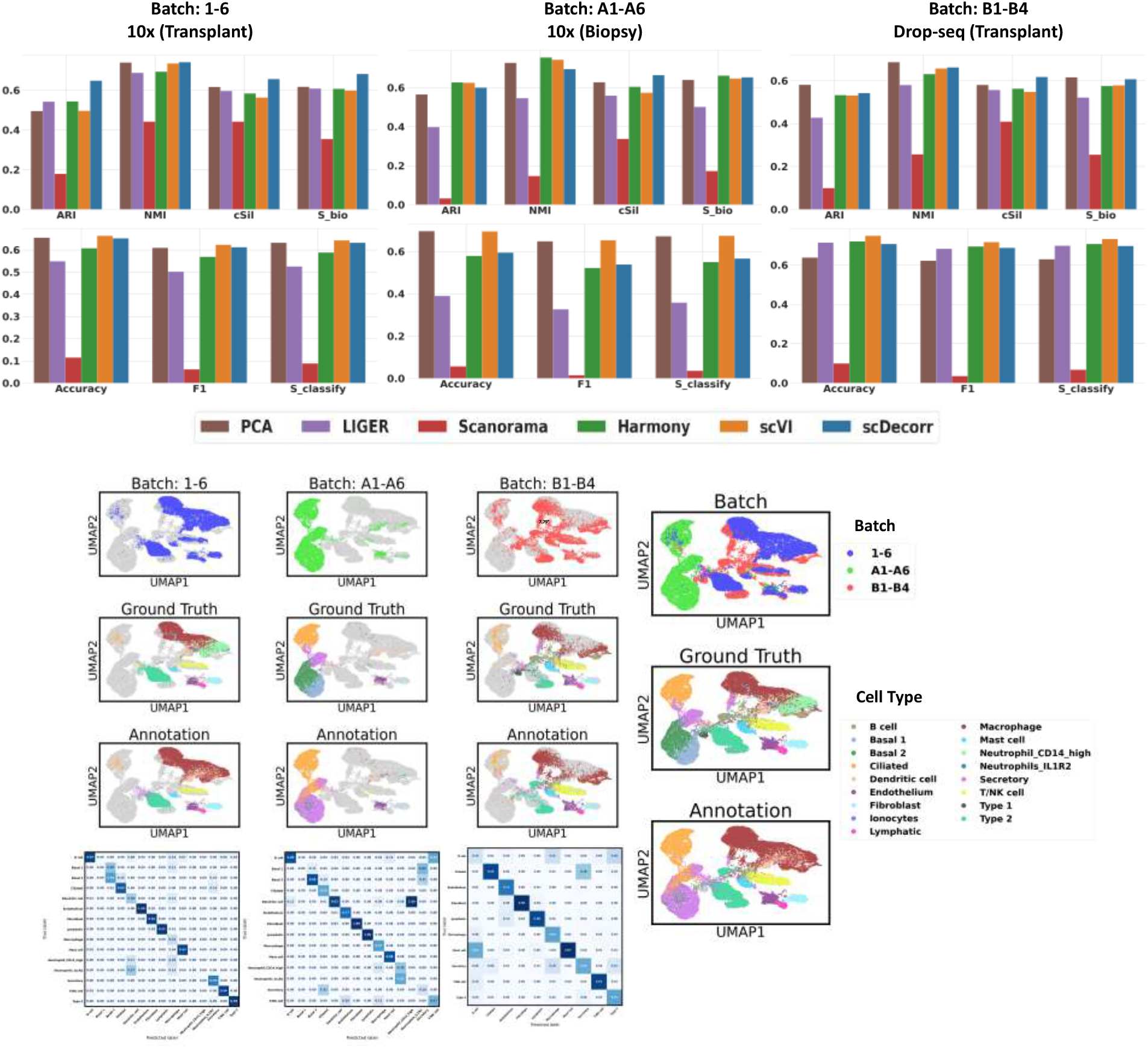
Human Lung data - Benchmark results of label transferring across studies. Here, two datasets act as reference while the third one acts as the query dataset with blinded cell-type labels (Table S4). An XGBoost model is trained on the reference dataset and cell-type classification task is performed for the left out query dataset. scDecorr shows comparable performance to other methods tested here. UMAPs show the batches, author’s annotations, and scDecorr predicted cell-types. Heatmaps show the overlap between predicted and author annotated cell-types.

We employ scDecorr to integrate the Human Lung dataset across its 16 donors. In Figure 2 (Figure 2), we showcase the data integration performance of scDecorr in comparison to other benchmark methods. Notably, on this dataset, scDecorr achieves a high biological conservation score (*S*_*bio*_) compared to alternative methods compared here. With an ARI (Adjusted Rand Index) of 0.636 and a cell-type silhouette score of 0.623, scDecorr slightly outperforms other methods. The next best performing method, scVI, achieves scores of 0.541 (ARI) and 0.543 (silhouette), falling slightly short of scDecorr’s performance in this dataset (Table S3).

However, it is noted that scDecorr’s batch correction scores are comparatively lower, particularly when compared to Harmony and LIGER (Table S3). This distinction is shown in the UMAP plots presented in Figure 2. Unlike Harmony or LIGER, scDecorr’s computed embeddings for cells from the 10x-Biopsy experiment (batches A1-A6) exhibit minimal overlap with embeddings from other experiments (Table S4). This outcome is desirable because these experiments have very few shared cell types with similar distributions. On the other hand, it could be that Harmony tends to over-correct the batches, leading to the merging of Basal 1, Basal 2, Secretory, and Type 2 cells. Additionally, Harmony struggles to accurately differentiate between Macrophage and Neutrophil-CD14-high cells. Similarly, LIGER also over-corrects the batches, merging Basal 1, Basal 2, Ciliated, and Type 2 cells.

From an overall perspective of data integration, scDecorr achieves a superior overall score (*S*_*overall*_) and outperforms other methods (Table S3). Subsequently, we utilize scDecorr for the task of reference to query cell-type label transferring. We annotate cells from each study (query) by using cells from the other studies as reference. For instance, while annotating 10x (Transplant) study, we use the 10x (Biopsy) and Drop-seq (Transplant) studies as references. The study-wise UMAPs and label transfer metrics barplots are shown in Figure 3.

When performing label transferring to the 10x (Transplant) and Drop-seq (Transplant) studies, scDecorr exhibits superior clustering (biological conservation) performance compared to all other methods (Table S4). Furthermore, scDecorr achieves competitive classification scores (Accuracy, F1) when compared to other methods for these two studies, while scVI emerges as the best-performing method. On the other hand, when label transferring to the 10x (Biopsy) study, scDecorr demonstrates competitive clustering performance, while scVI emerges as the best-performing method in terms of classification scores for this particular study (Table S4). In this dataset, the annotation performance of scDecorr, although not as good as scVI, remains on par with Harmony. It is important to note that the 10x (Biopsy) study exhibits minimal overlap in cell types with the other studies. Therefore, the clustering scores provide a more comprehensive evaluation of the label transferring performance across all cell types in this study. Interestingly, PCA also achieves high clustering and classification scores across all the query studies.

##### Integration of Human Kidney datasets

We integrate a total of 224,696 human kidney cells from three different datasets: KPMP (KP), BUKMAP (BU), and CKD (CD). Here, KPMP has 105,401 cells, BUKMAP has 67,446 cells, and CKD dataset has 51,849 cells. There are 34 distinct cell types in the combined datasets. Here, throughout our experiments, we treat each dataset as a separate batch. The KPMP and BUKMAP datasets share 16 cell types, which are present in both batches. However, the CKD dataset, which has the smallest number of cells in the overall dataset, contains 29 unique cell types. It’s worth noting that the cell type naming and annotation resolution in CKD differ from the other batches. Among the 29 ontologies in the CKD dataset, 11 are shared with KPMP and BUKMAP, while the remaining 18 ontologies are exclusive to CKD. Additionally, the majority of cell type annotations in CKD are not present in the other batches. This skewed distribution of cell types in the CKD study adds complexity to the overall use case.

The data integration performance of scDecorr, compared to other benchmark methods, is illustrated in Figure 4 (Table S5, Table S6). The barplots show that scDecorr outperforms other methods in terms of biological conservation and overall score. While LIGER and Harmony excel in batch mixing, scDecorr’s batch correction performance is slightly lower than other methods. However, when considering the overall data integration performance (*S*_*overall*_), scDecorr is comparable to other methods. Interestingly, PCA also achieves significantly high overall performance. To understand the relatively lower batch mixing scores of scDecorr, we analyzed the label transfer i.e. predicted cell-types by scDecorr (Figure 5). As shown, the low *S*_*batch*_ score of scDecorr could be due to the fact that the cell type distributions in the CKD study differ significantly from the other two batches.

**Figure 4.**
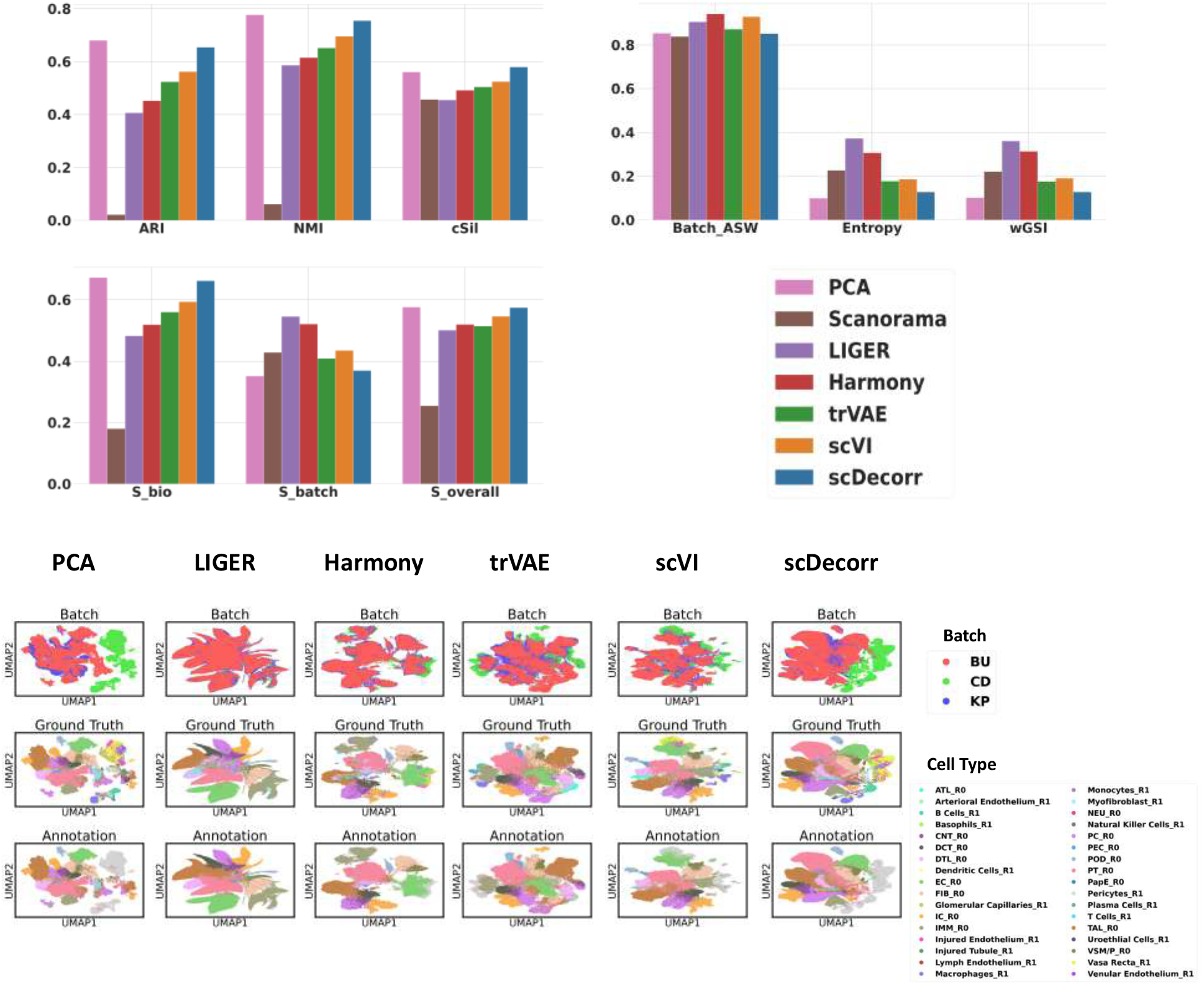
Human Kidney: Data Integration Benchmark Results. We applied scDecorr to integrate 224,696 Human kidney cells from the KPMP, BUKMAP and CKD datasets. scDecorr has the highest *S*_*bio*_ and *S*_*overall*_ scores (Table S5). 2D representation of batch, author annotated cell-types and overall predicted cell-types is shown as UMAP.

**Figure 5.**
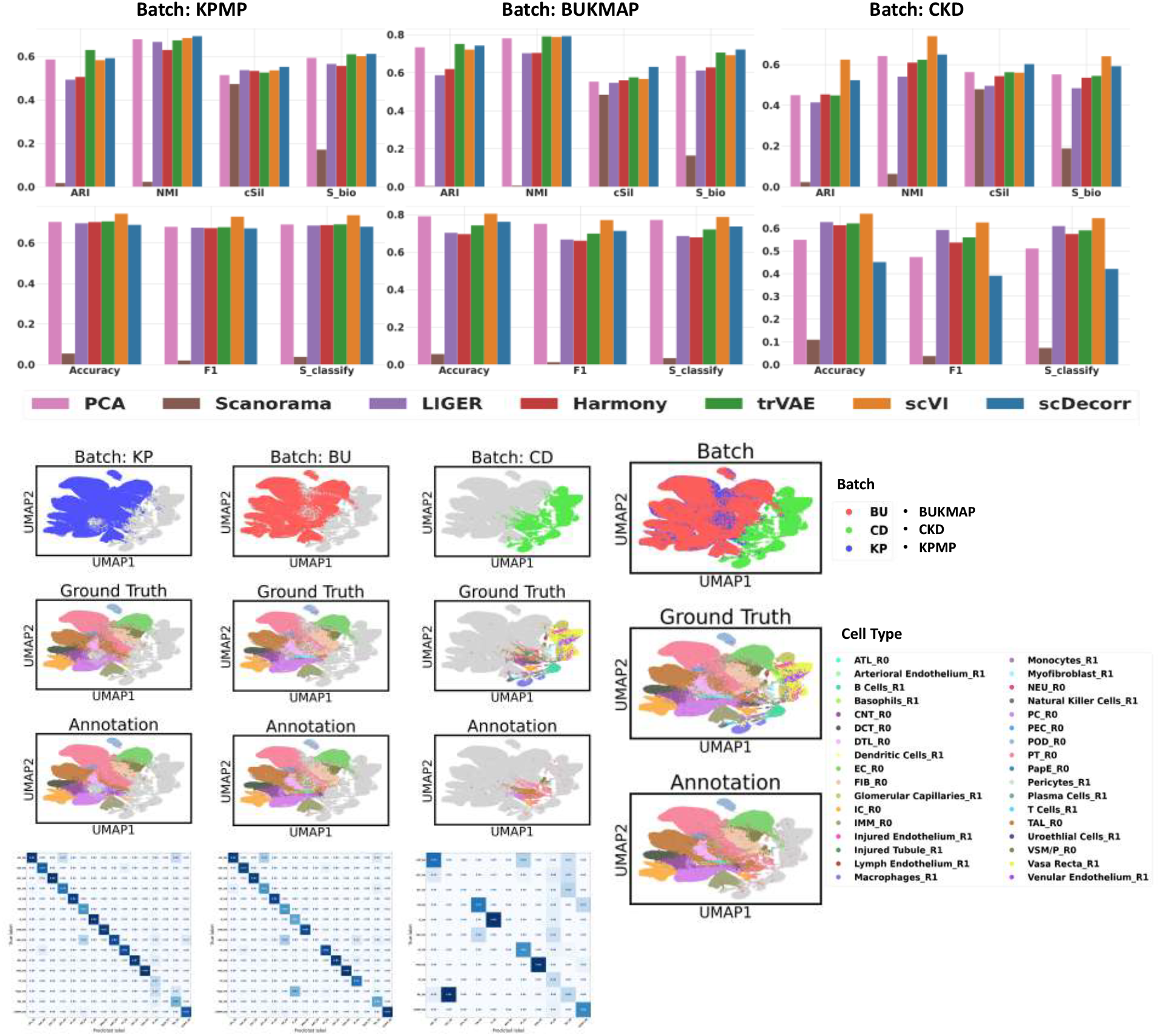
Human Kidney: Benchmark results for label transferring across datasets. Barplots show the Accuracy, F1 and *S*_*classify*_ scores for different methods. scDecorr shows high scores for the KPMP and BUKMAP datasets while scVI works well for all three datasets ((Table S6).

The majority of cell types in the CKD study, namely Vasa Recta, Glomerular Capillaries, Venular Endothelium, Injured Endothelium, B Cells, and Macrophages, comprise nearly 70% of the cells in the CKD study but are absent in the other batches. Examining the CKD UMAP plot in Figure 5, we can observe that the regions of CKD that do not overlap with KPMP and BUKMAP predominantly consist of these six majority cell types, along with a few other types that are unique to CKD. This outcome is desirable since we aim to avoid mixing different cell ontologies from different batches. Additionally, scDecorr successfully integrates shared cell types of CKD, such as PT, TAL, PC, and DTL, with corresponding cell ontologies from the other batches. However, we also notice that scDecorr does not integrate some shared cell types of CKD, like IC and DCT, effectively with the KPMP and BUKMAP batches.

We further evaluated the label transferring performance of scDecorr by treating each dataset as a query while using the others as references. For example, when annotating the KPMP study, we utilized the BUKMAP and CKD studies as references. The study-wise UMAPs and barplots displaying label transfer metrics are presented in Figure 5. When transferring labels to the KPMP and BUKMAP studies, scDecorr exhibits superior clustering performance compared to other methods, indicating better preservation of biological information (Table S6). Additionally, scDecorr achieves competitive classification scores, although scVI stands out as the top-performing method for these two studies. On the other hand, when transferring labels to the CKD study, scDecorr demonstrates competitive clustering performance, while scVI emerges as the best-performing method in terms of clustering scores. However, when it comes to classification performance on CKD, scDecorr performs noticeably worse than other methods, with scVI being the best method. It is important to note that the CKD study displays minimal overlap in cell types with the other datasets. Consequently, the clustering scores provide a more comprehensive evaluation of the label transferring performance across all cell types in this dataset. Interestingly, PCA achieves high clustering and classification scores across all the query studies.

Figure 5 and Figure 6 show the prediction accuracy and UMAPs for the label transfer based predicted cell-types for the five other benchmark methods used in the study. Further, Figure 4 shows a low *S*_*batch*_ score for PCA based integration, this is due to a prominent batch effect between CKD and the other batches that can not be corrected by PCA alone 6A. On the other hand, *S*_*batch*_ score for Harmany is high as shown in Figure 4, the corresponding UMAP, is shown in Figure 6B. However, this integration comes at the expense of over-correction and a sacrifice of biological preservation. trVAE has a low score for both batch correction and biological conservation (Table S5 and S6), as indicated by the evaluation metrics presented in Figure 4. The illustrative UMAP for trVAE based integration is shown in Figure 6C. scVI, as depicted in the UMAP plots in Figure 6D, successfully mixes the batches while preserving biological conservation to a reasonable extent. However, similar to Harmony, scVI also attempts to merge unique CKD cell types, such as Vasa Recta and Venular Endothelium, with EC cells from KPMP and BUKMAP. It also aligns Injured-Tubule cells of CKD with PT and PC cells from KPMP and BUKMAP. While LIGER achieves perfect batch mixing, it fails to preserve biological information specific to the CKD batch. As shown in Figure 5 and illustrated as UMAP in Figure 6E, it merges all the unique cell types of CKD with different cell types from KPMP and BUKMAP.

**Figure 6.**
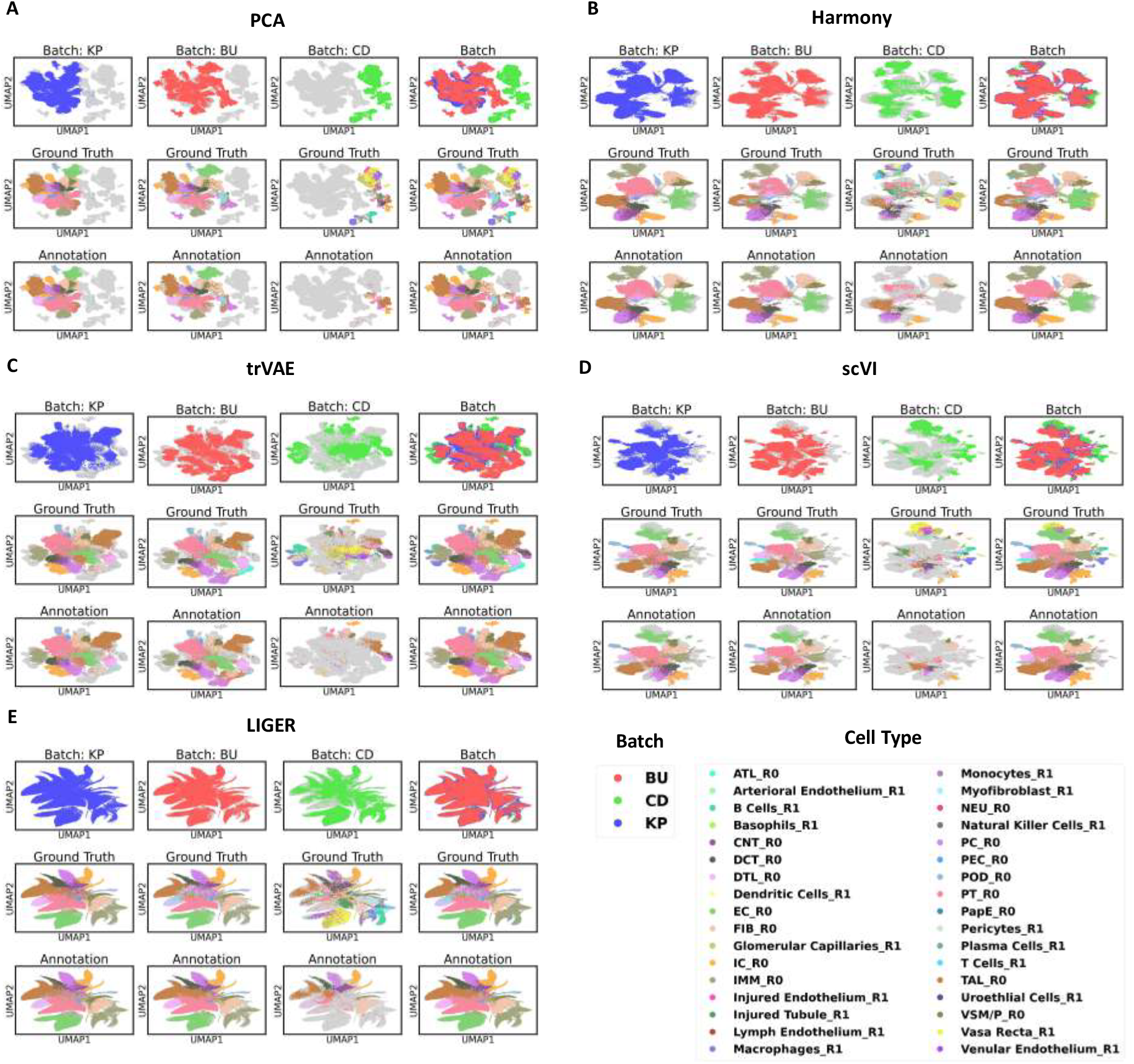
Human Kidney: UMAP plots of all methods for label transferring across datasets. Here cell-types were predicted in a leave one out manner, where two datasets were used as reference and cell-types were predicted for the blinded query dataset.

#### 3.5.2 Case Study 2: Benchmarking scDecorr on datasets with significant number of shared cell types across batches Integrating Cross-tissue Immune Cells across 10x Chemistries

We integrate the “T and & innate lymphoid cells” subset of the Cross-tissue Immune Cell Atlas[38] across different chemistries using scDecorr. This specific subset comprises 216,611 cells that were derived from 17 different organs including Lung, Lymph Nodes, Liver, Spleen, etc. The cells were sequenced using three different chemistries: 5’ v1, 5’ v2, and 3’ provided by 10x Genomics technology. Each chemistry accounts for 41,715, 61,559, and 113,337 single cells, respectively. The dataset encompasses 18 types of T and innate lymphoid cells originating from 17 distinct organs. Notably, all cell types are present in all three chemistries. However, when examining the raw data, there is a notable batch effect between the chemistries, which is clearly visible in the UMAP plot of PCA features depicted in Figure 7. Consequently, to address this issue, we integrated the dataset using the “Chemistry” variable as the batch key.

**Figure 7.**
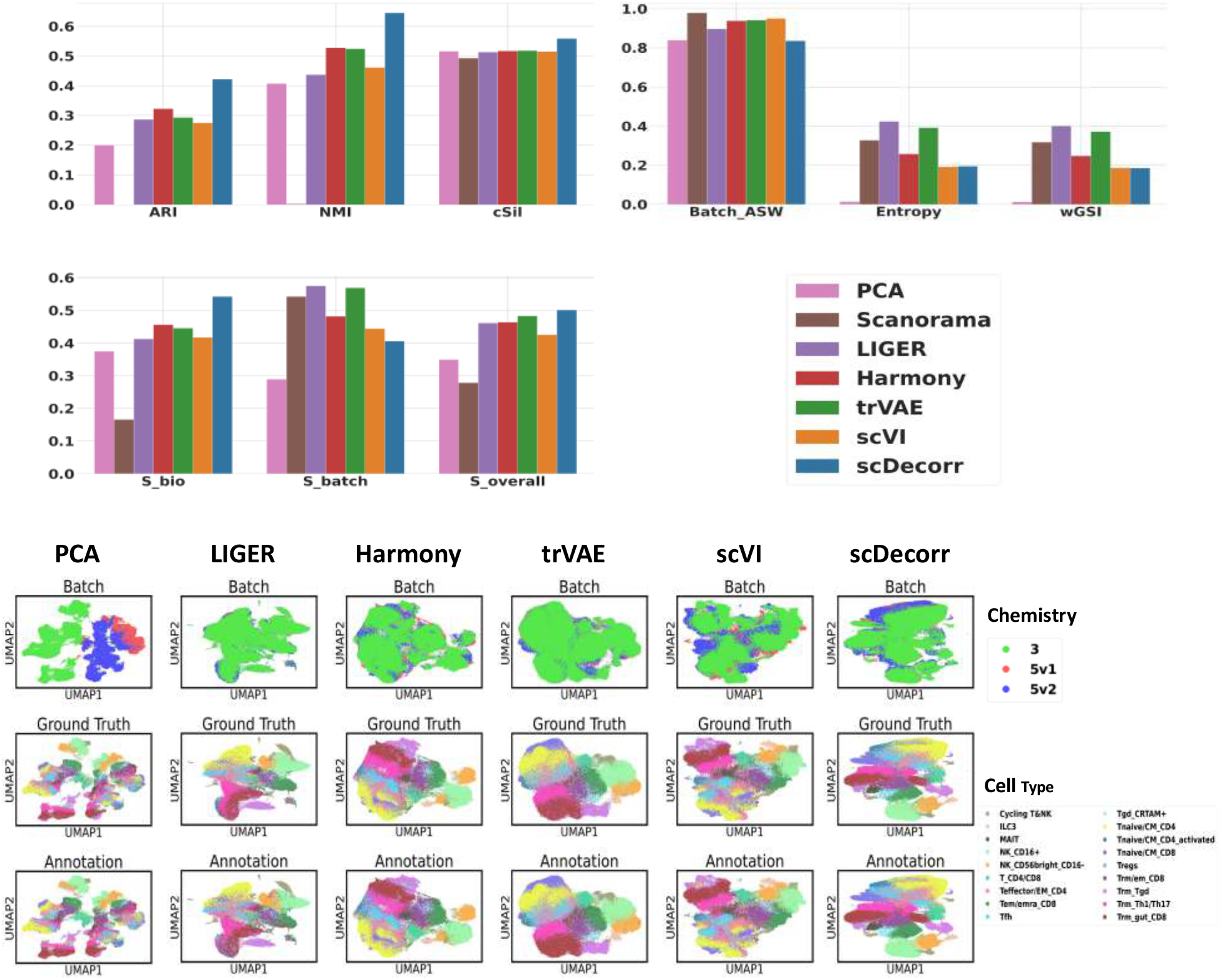
Cross-tissue Immune atlas: Benchmark Results of Data Integration across Chemistries. scDecorr is used to integrate different immune datasets with chemistries available in 10X sequencing. scDecorr achieves the highest overall *S*_*bio*_ and *S*_*overall*_ score for datasets tested here (Table S7).

Figure 7 (Table S7 and S8) shows the performance of scDecorr compared to other benchmark methods in terms of data integration. scDecorr surpasses other methods by a significant margin in terms of biological conservation performance (*S*_*bio*_). Here, scDecorr exhibits approximately a 20% improvement in the *S*_*bio*_ score compared to the second-best performing models, namely Harmony and trVAE. The NMI (Normalized Mutual Information) score for scDecorr is 0.645, while Harmony and trVAE achieve scores of 0.528 and 0.525, respectively.

The batch correction barplots in Figure 7 show that scDecorr’s batch mixing scores are notably lower than those of the other methods. However, we believe that scDecorr effectively mixes the three batches, and unlike Harmony or trVAE, scDecorr avoids over-correction of the batches. This observation is supported by the number of cell clusters identified by the Leiden clustering algorithm. Out of the 18 ground truth cell types, scDecorr successfully identifies 15 cell types, while Harmony, trVAE, and scVI features identify only 9, 5, and 8 cell types, respectively. In terms of overall data integration (*S*_*overall*_), scDecorr outperforms all the other methods (Figure 8). The cell-type clusters obtained from scDecorr are clearly distinguishable from one another. Conversely, the cell embeddings produced by Harmony and trVAE fail to form distinct cell type clusters. These methods tend to excessively correct the batches, causing unrelated cell types to group closely together in the representation space, thus resulting in sub-optimal clustering performance (biological conservation), which is also reflected in the *S*_*bio*_ scores (Table S7 and S8).

**Figure 8.**
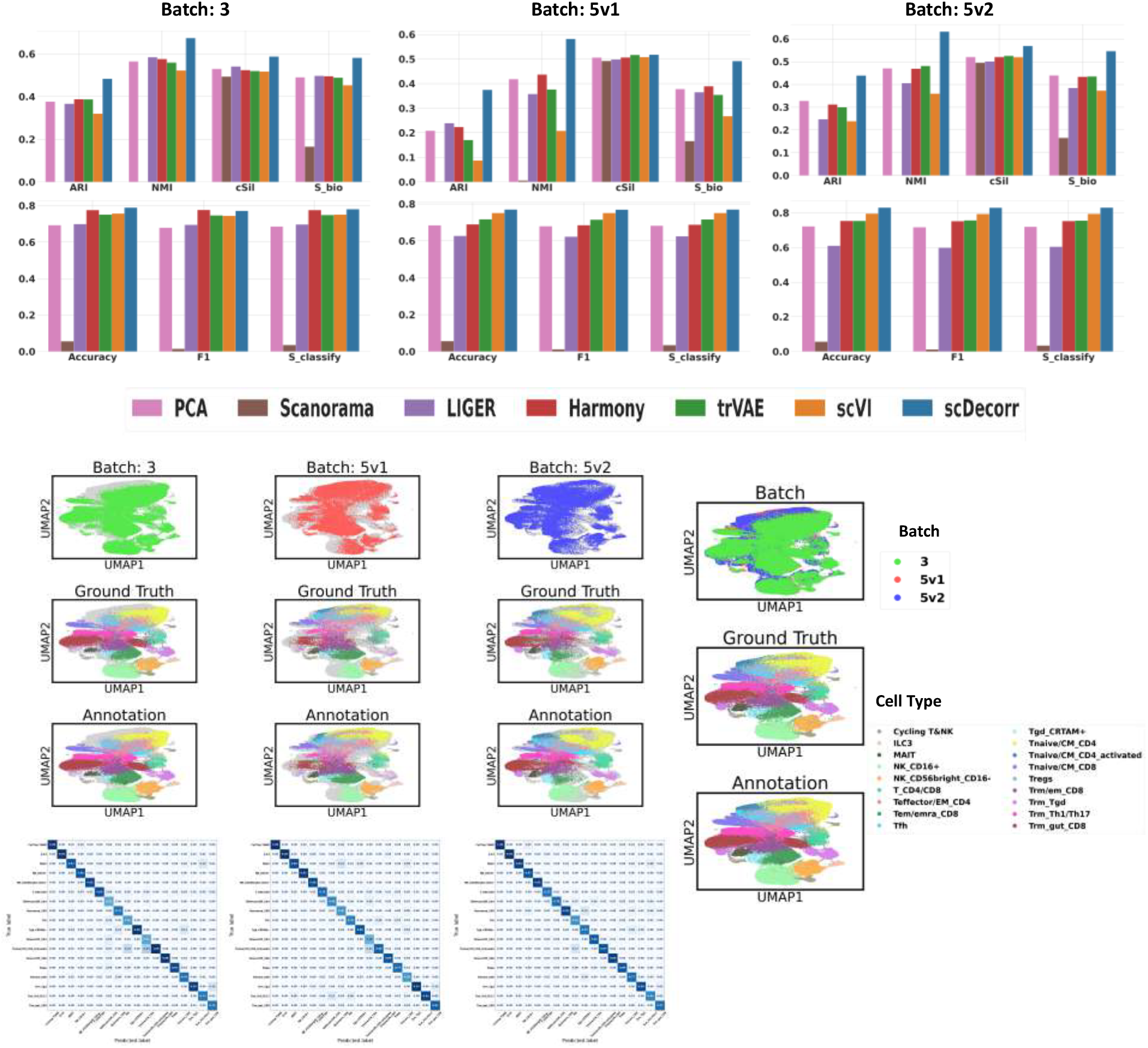
Cross-tissue Immune: Benchmark results of label transferring across chemistries. Barplots show accurcy, F1 and *S*_*classify*_ of the different methods tested here. Integrated datasets are shown as UMAPs. Predicted and observed cell-types obtained using scDecorr show that scDecorr’s embedding can be used for optimal clustering and automated cell-type annotation (Table S8).

Next, we analyzed the Chemistry-wise label transferring performance of scDecorr(Figure 8). The clustering metrics show that scDecorr significantly outperforms all other methods across all chemistries. Notably, on the 5’ v1 batch, scDecorr exhibits approximately 60% improvement in Adjusted Rand Index (ARI) and around 30% improvement in Normalized Mutual Information (NMI) compared to the alternative methods. These results highlight scDecorr’s exceptional ability to preserve the biological signal consistently across all chemistries. Moreover, Figure 8 reveals that scDecorr surpasses all other methods in classification performance (accuracy and F1 score) on the 5’ v1 and 5’ v2 chemistries. Additionally, scDecorr achieves competitive performance on the 3’ chemistry batch. The heatmaps show the overlap between predicted and observed cell-type labels and illustrate scDecorr’s effectiveness in allowing shared cell types from all three batches to cluster together without the batches being excessively corrected. In summary, our analysis (Figure 8) shows scDecorr’s higher performance in label transferring across different chemistries in the cross-tissue immune dataset. Here, scDecoor outperforms other methods by in terms of clustering and classification metrics. Additionally, scDecorr successfully integrates the chemistries in a manner that facilitates the clustering of shared cell types without over-correction of the batches.

In our final analysis, we utilize the integrated representations generated by scDecorr to assess the performance of inter-organ label transfer. To this end, we treat cells originating from one organ as queries and the remaining organs as references. The organ-wise annotation UMAPs produced by scDecorr are illustrated in Figure 9. Furthermore, we present the label transfer benchmark plots in Figure 10, allowing us to observe the performance of scDecorr in comparison to other methods. Upon examination of the figures, it becomes evident that scDecorr outperforms all other methods for every organ considered, displaying superior clustering and classification performance across the board. The results emphasize the effectiveness of scDecorr in accurately transferring labels between different organs.

**Figure 9.**
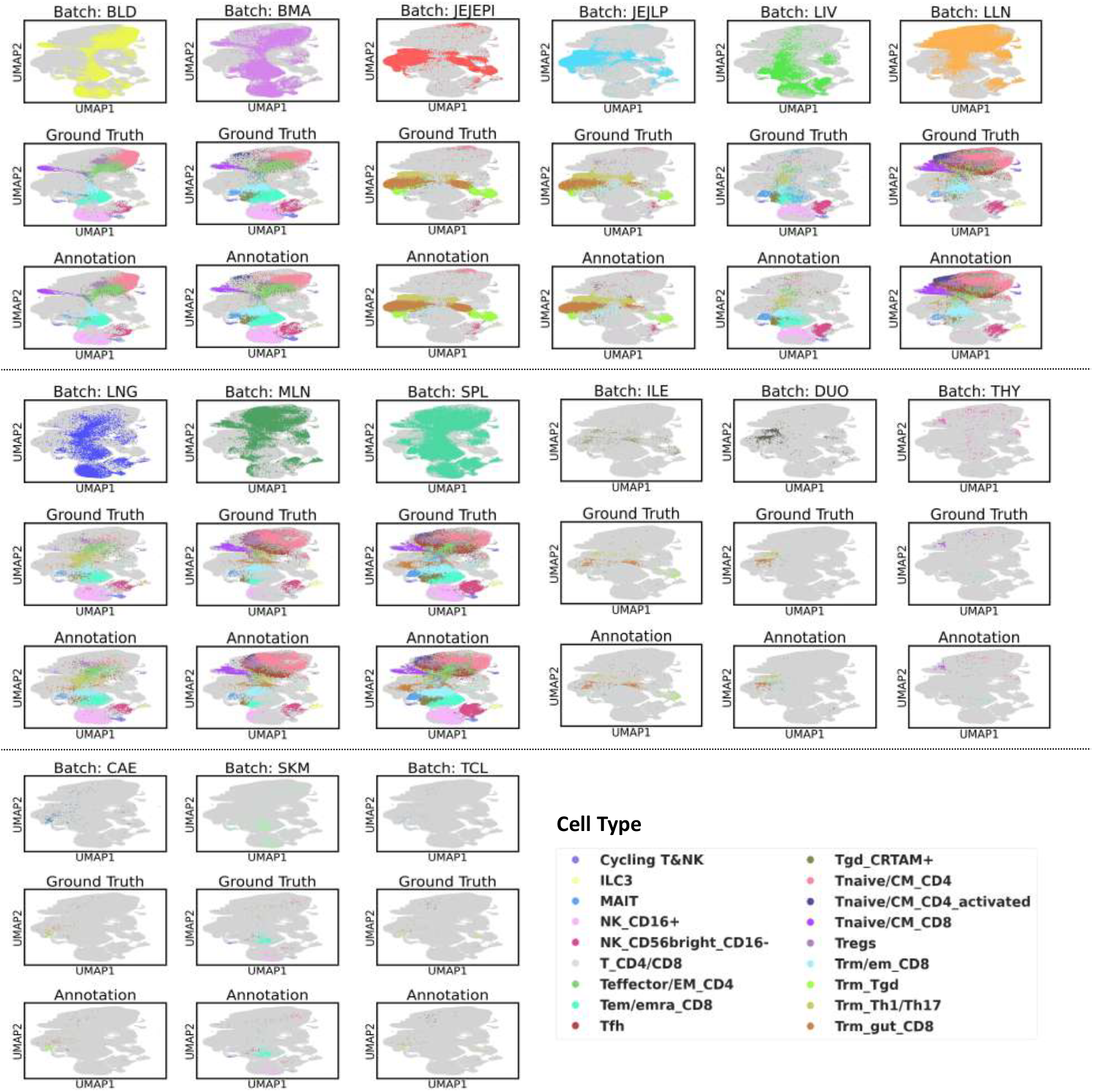
Cross-tissue Immune: UMAP plots of inter-organ label transfer using scDecorr for each organ in the dataset. Ground Truth refers to the annotations provided by the authors in their original study.

**Figure 10.**
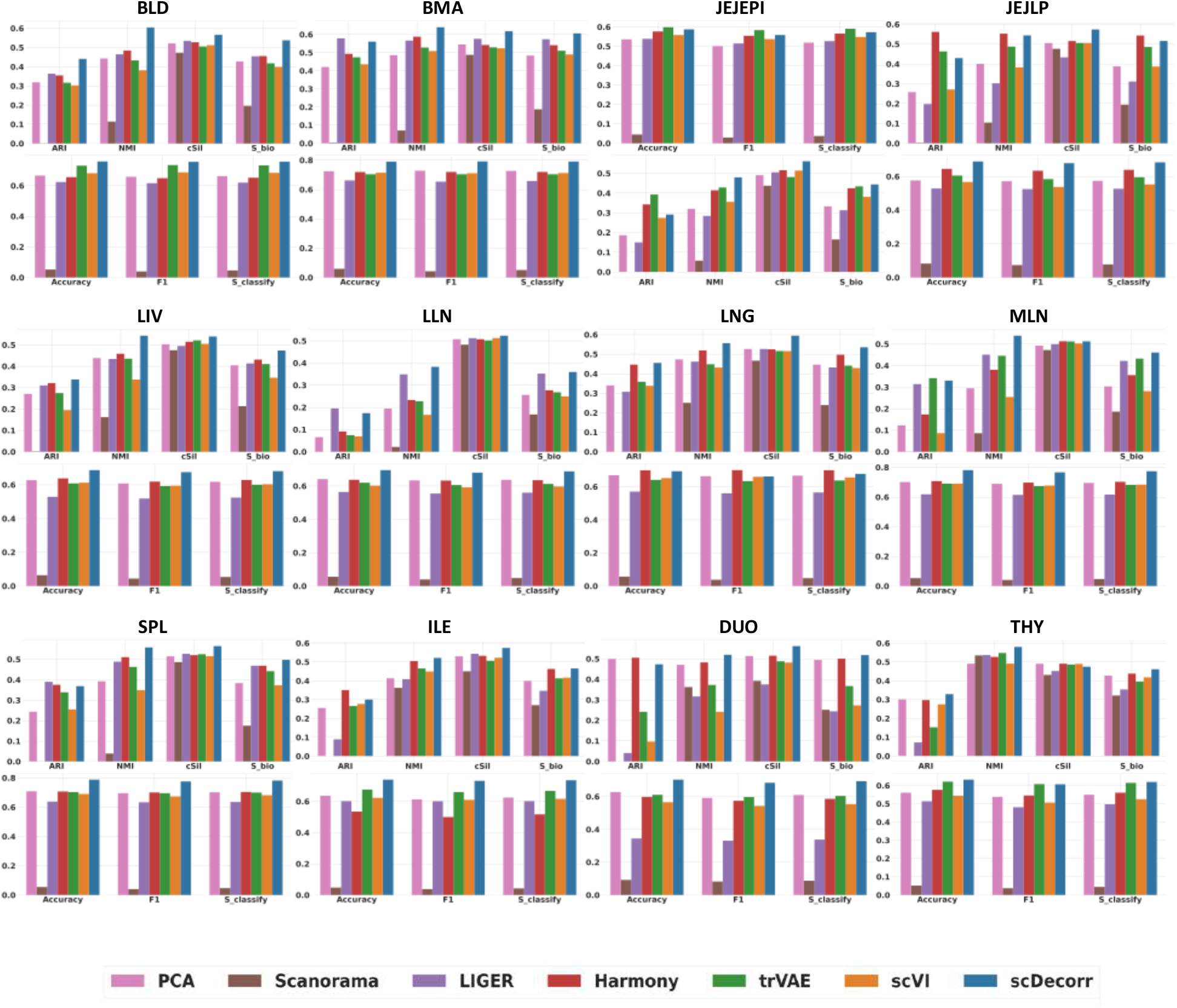
Cross-tissue Immune: Benchmark results of inter-organ label transfer. For each method, prediction accuracy measures (Accuracy, F1 and *S*_*classify*_), clustering metrices such as NMI and ARI and bio conservation scores *S*_*bio*_ are provided for each organ.

##### Integrating Tabula Muris across technologies

The Tabula Muris dataset[3], comprises 67,354 single cells sequenced from 24 distinct mouse organs using Droplet and FACS technologies. The Droplet and FACS technologies contributed 47,664 and 19,690 cells, respectively. Within the dataset, there are 28 unique cell types, with most cell types being shared between the two technologies. However, PCA based UMAP plot shows (Figure 11) a noticeable batch effect between the droplet and FACS technologies. Consequently, we address this issue by integrating the dataset, utilizing technology as the batch key. The barplots in Figure 11 demonstrate that scDecorr exhibits competitive biological conservation scores when compared to other benchmark methods. Specifically, scDecorr achieves an ARI of 0.68, NMI of 0.80, and cell-type silhouette (cSil) score of 0.59. Moreover, scDecorr achieves a high kBET score of 0.856, indicating its successful integration of the technologies. Overall, scDecorr’s data integration performance (represented by *S*_*overall*_) is comparable to that of other methods (Table S9).

**Figure 11.**
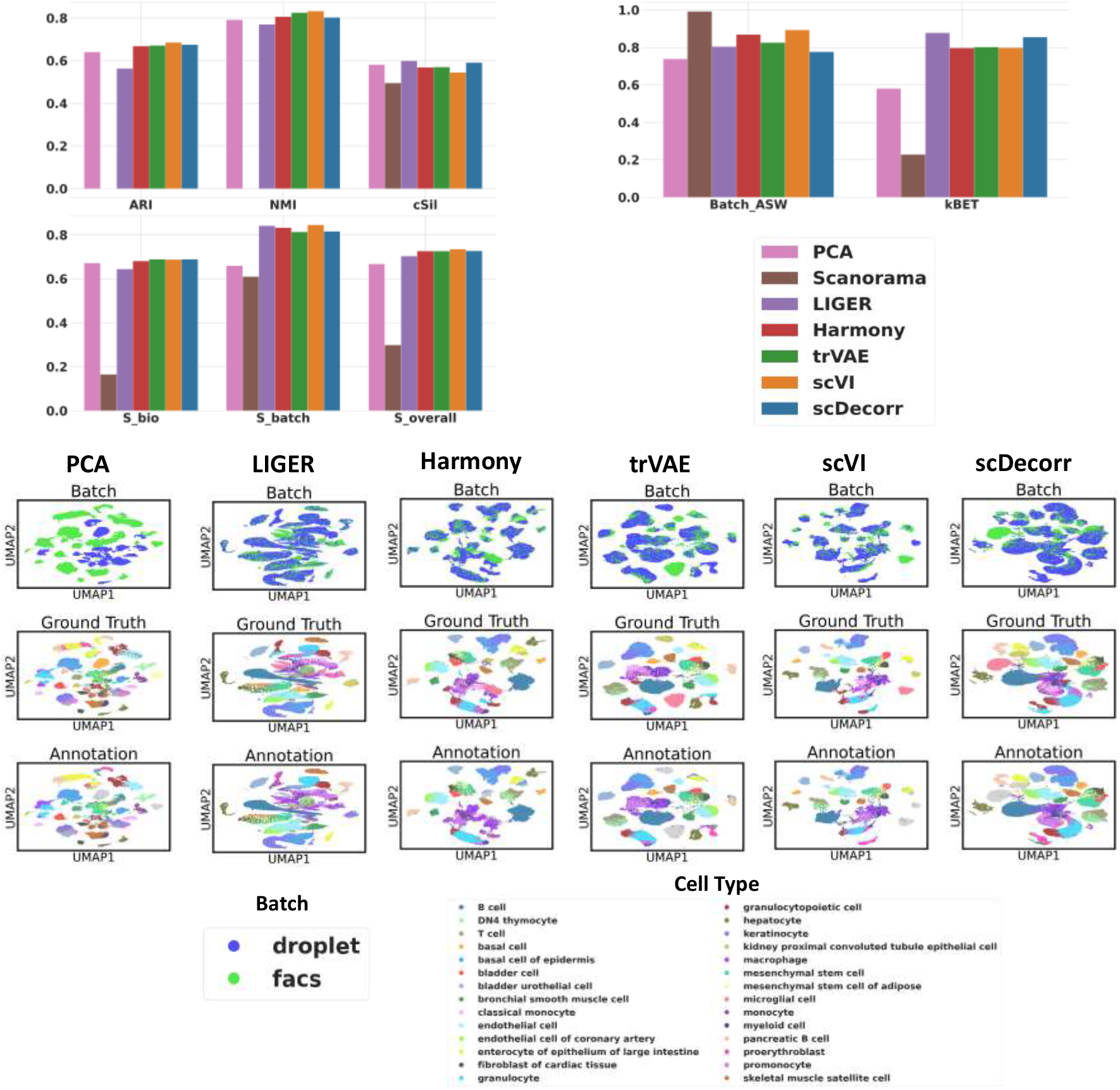
Tabula Muris: Benchmark Results of Data Integration across Technologies. Barplots show 8 different metrices of comparing 7 different methods on the task of data integration. UMAP plots show batch, author’s annotation and scDecorr predict cell-type labels (Table S9).

Next, we evaluate scDecorr’s performance in terms of technology-wise label annotation by transferring labels between Droplet and FACS (Table S10). Figure 12 (Table S9, S10) shows the annotation and biological conservation results of the query batches. In both technology batches, scDecorr demonstrates competitive balanced accuracy and F1 scores. The clustering metrics barplots in Figure 12 further illustrate scDecorr’s ability to preserve the inherent biological signal of each batch. In our final analysis, we employ the integrated representations generated by scDecorr to assess the performance of inter-organ label transfer. Here, cells from one organ serve as query, while the remaining organs act as references. Figure 13 showcases the organ-wise annotation UMAPs produced by scDecorr, while Figure 14 presents the label transfer benchmark plots, allowing for a comparison with other methods. Figure 14 shows that scDecorr achieves competitive label transfer performance for all organs. These results showcase scDecorr’s effectiveness in accurately transferring labels between different organs.

**Figure 12.**
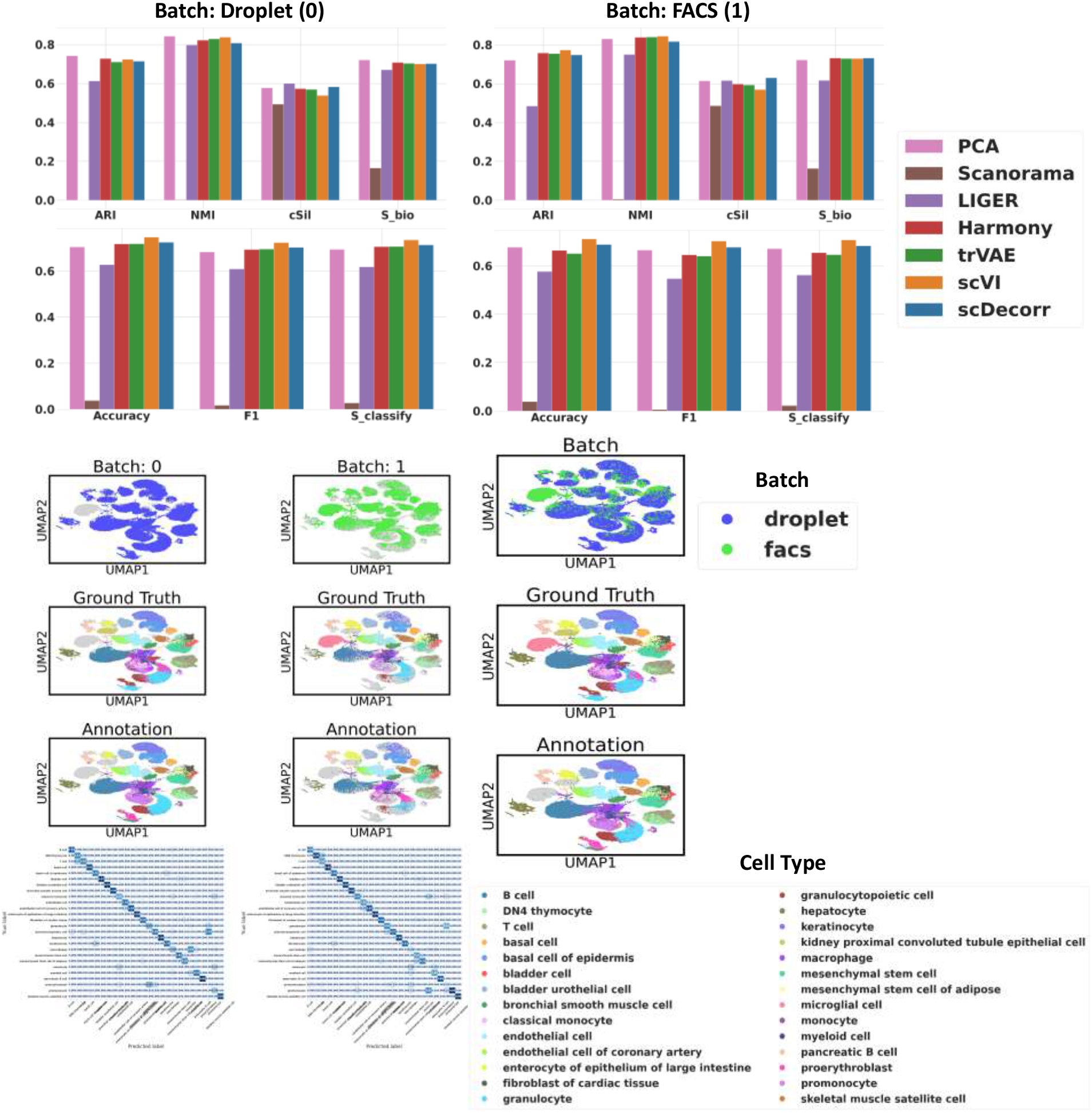
Tabula Muris: Benchmark results of label transferring across technologies. scDecorr can be used to integrate datasets from droplets and FACS based technologies (Table S10).

**Figure 13.**
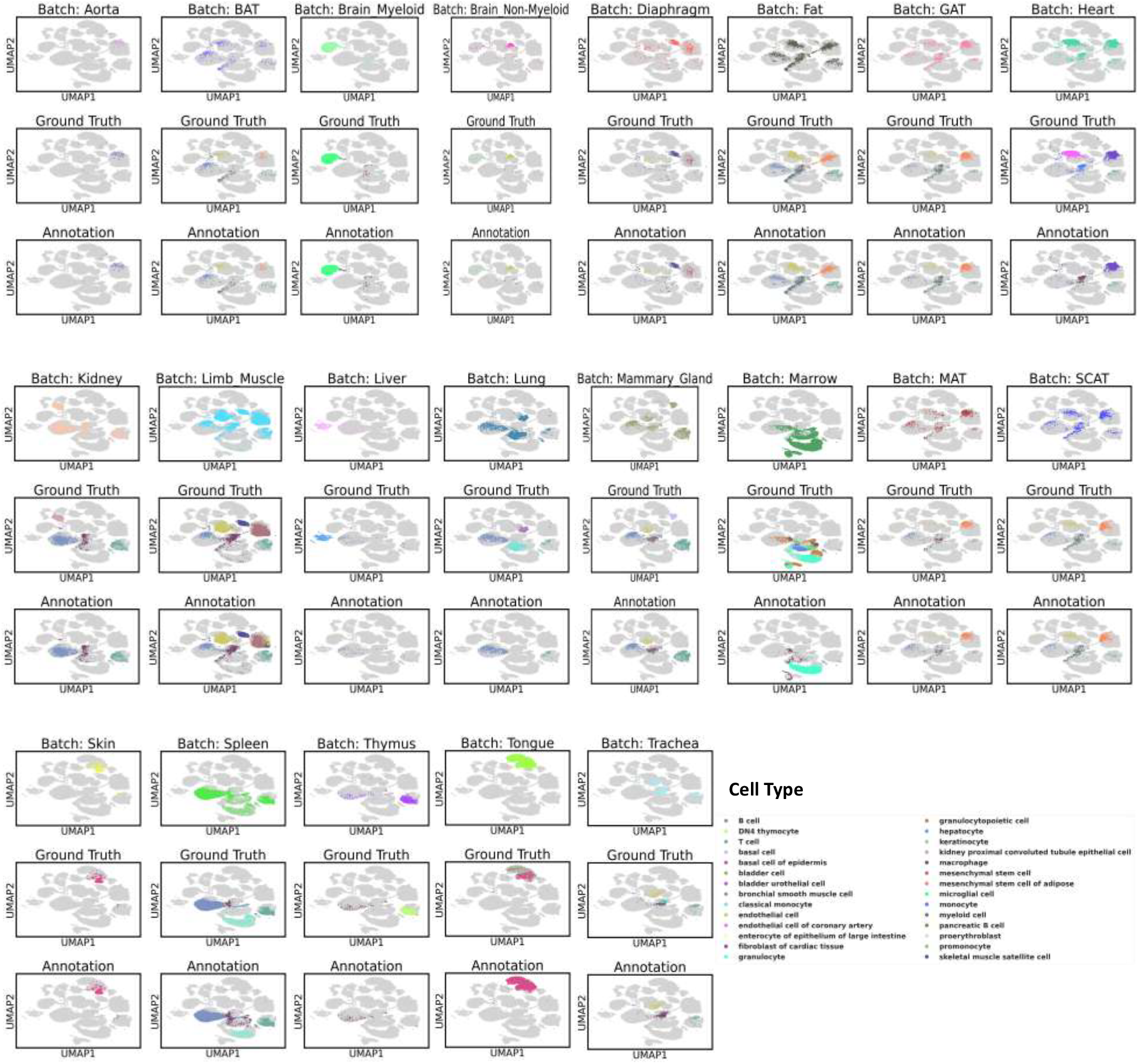
Tabula Muris: UMAP plots of inter-organ label transfer using scDecorr. Here, scDecorr’s embedding was used to traing an XGBoost model using cell-labels of all except the query organ, which was used in a blinded manner for predicting the cell-types.

**Figure 14.**
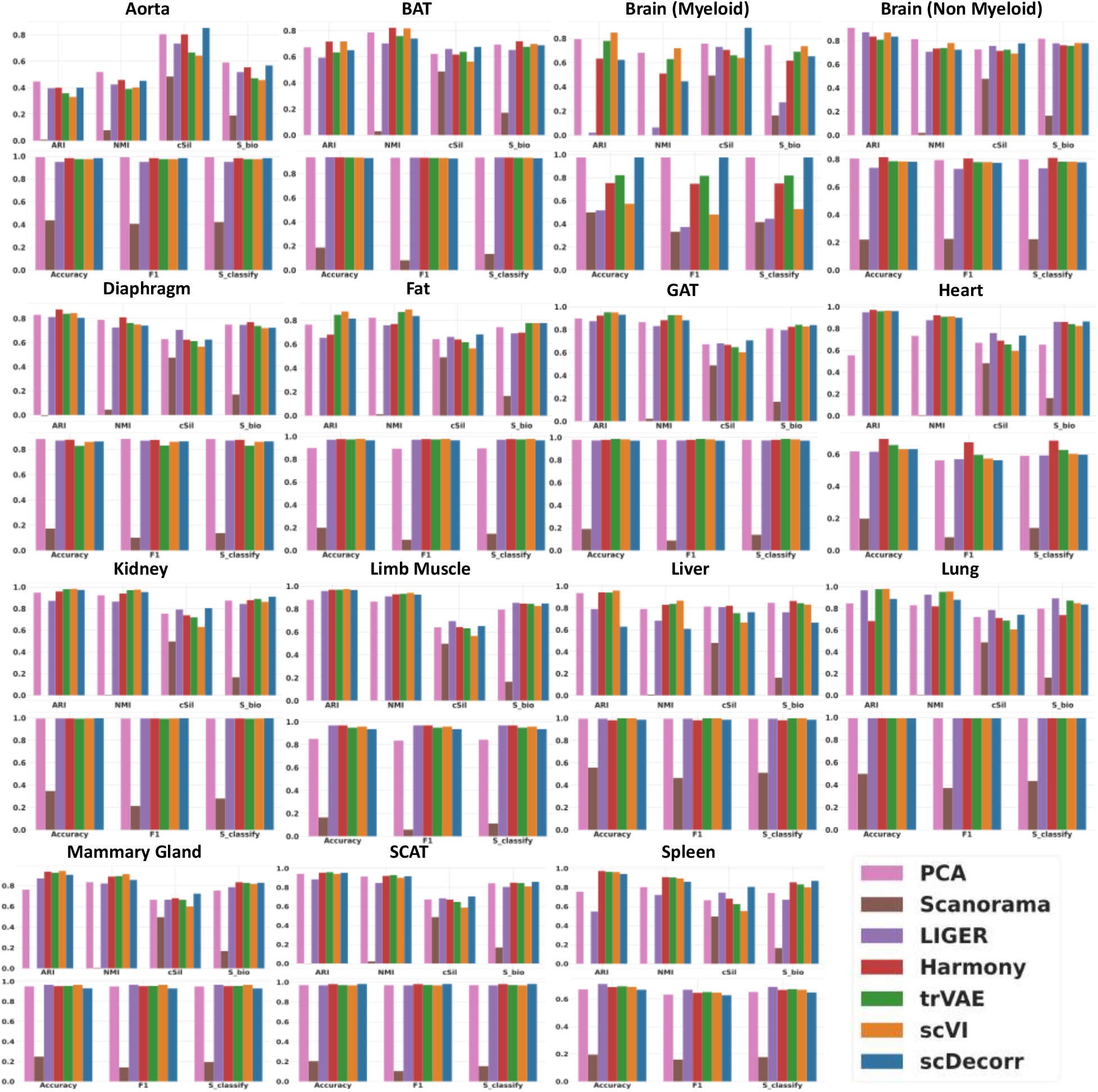
Tabula Muris: Benchmark results of inter-organ label transfer. Prediction and cluster accurcy metrices for different methods used in this study show the utility of embeddings generated by scDecorr for downstream tasks.

## 4 Discussion

By treating each cell as a distinct entity, scDecorr employs a strategy that maximizes alignment between the joint embeddings of gene profiles while simultaneously decorrelating their components. This approach achieves two key objectives - firstly, it maximizes similarity among the cell embeddings of distorted profiles ensuring that the representations become invariant to the applied distortion. And secondly, by decorrelating the components of the embedding vectors, scDecorr captures a highly variable representation space in a self-supervised manner, without relying on negative examples.

In the process of integrating unannotated cells from multiple domains (such as sources, experiments, or batches), scDecorr employs domain adaptation to learn domain-invariant cell representations. It accomplishes this by independently learning cell representations across domains through random sampling of instances from each domain. Unlike most single-cell data integration methods, scDecorr does not explicitly perform batch correction. Instead, it aims to capture a shared latent space for all batches by simultaneously learning the optimal latent space for each batch in a self-supervised manner. This implicit batch mixing process allows scDecorr to better preserve biological information, as evidenced by improved clustering scores in the latent space. Moreover, since there is no explicit batch correction, scDecorr avoids over-correction of the batches.

To assess the data integration performance of scDecorr, we apply it to datasets with varying degrees of shared cell types between batches, considering diverse integration conditions such as different donors, species, studies, chemistries, and platforms. The results demonstrate that scDecorr successfully integrates the batches in a manner that facilitates the clustering of shared cell types without over-correction. Additionally, the representations computed by scDecorr exhibit robustness in label transfer tasks, allowing for effective transfer of labels from reference to query datasets.

## Acknowledgments

SH and RS were supported in part by the Leducq Early Career Investigator Seed Grant Award in the IMMUNO-FIB HF Network, CRU344, RWTH START and Novo Nordisk STAR grant.

## Supplementary File

scDecorr

